# Strain diversity of plant-associated *Lactiplantibacillus plantarum*

**DOI:** 10.1101/2021.03.28.437403

**Authors:** Annabelle O. Yu, Elissa A. Goldman, Jason T. Brooks, Benjamin L. Golomb, Irene S. Yim, Velitchka Gotcheva, Angel Angelov, Eun Bae Kim, Maria L. Marco

**Author notes:** Correspondence: Maria L. Marco, One Shields Avenue, University of California, Davis, Davis, CA 95616.

## Abstract

The intraspecific phenotypic and genetic diversity of *Lactiplantibacillus plantarum* (formerly *Lactobacillus plantarum*) was examined for five strains isolated from fermented olives and eight strains from cactus fruit, fermented tomatoes, teff injera, wheat boza, and wheat sourdough starter sources. Carbohydrate utilization and stress tolerance characteristics showed that the olive isolates grew more robustly in galactose and raffinose, showed higher tolerance to 12% v/v EtOH, and exhibited a greater capacity to inhibit an olive spoilage strain of *Saccharomyces cerevisiae* than *L. plantarum* from the other plant sources. Certain traits were variable between fermented olive isolates such as the capacity for biofilm formation and survival at pH 2 or 50 °C. By comparison, all *L. plantarum* from fruit sources grew better at a pH of 3.5 than the strains from fermented grains. Multi-locus sequence typing and genome sequencing indicated that strains from the same source type tended to be genetically related. Comparative genomics was unable to resolve strain differences, with the exception of the most phenotypically impaired and robust isolates. The findings show that *L. plantarum* is adapted for growth on specific plants or plant food types, but that intraspecific variation may be important for ecological fitness of *L. plantarum* within individual habitats.

## Introduction

Certain LAB required for food fermentations are recognized for their genetic and phenotypic diversity and have been classified as “nomadic” or “generalist” because of their broad habitat range (Duar *et al*., 2017; Choi *et al*., 2018; Yu *et al*., 2020). *Lactiplantibacillus plantarum* (formerly *Lactobacillus plantarum* (Zheng *et al*., 2020)) is included among those nomadic LAB (Duar *et al*., 2017) and is known for its significant intraspecific versatility (Molenaar *et al*., 2005; Siezen *et al*., 2010; Martino *et al*., 2016). *L. plantarum* is frequently isolated from fresh and fermented plant, meat, and dairy foods and is an inhabitant of the gastrointestinal and vaginal tracts of humans and animals (Delgado *et al*., 2005; Aquilanti *et al*., 2007; Di Cagno *et al*., 2008; Yang *et al*., 2010; Ciocia *et al*., 2013; Jose *et al*., 2015; Zago *et al*., 2017; Parichehreh *et al*., 2018; Barache *et al*., 2020). This species is essential for the production of numerous fermented foods (e.g., fermented olives, sauerkraut, salami, and sourdough), and certain strains are effective probiotics (Marco, 2010; Seddik *et al*., 2017; Crakes *et al*., 2019). Consistent with its host and environmental range, *L. plantarum* strains have larger genomes compared with LAB with narrow host ranges and also carry strain-specific genes, often located on lifestyle adaptation islands (Molenaar *et al*., 2005; Sun *et al*., 2015; Zheng *et al*., 2015; Duar *et al*., 2017; Salvetti *et al*., 2018;).

Despite the robust growth of *L. plantarum* in different host-associated and food environments, *L. plantarum* genomes and cell properties have thus far shown limited correlations with isolation source across disparate habitats (Siezen *et al*., 2010; Martino *et al*., 2016). These findings indicate that either intraspecific variation of *L. plantarum* within individual sources is fortuitous and members of this species have not evolved for growth in specific habitats (Martino *et al*., 2016), or that this observed variation is the result of adaptive evolution of the *L. plantarum* species within certain habitats with the outcome of maximizing co-occurrence by niche complementarity (Bolnick *et al*., 2011; Ehlers *et al*., 2016).

To begin to address these two hypotheses, we examined the intraspecies variation of a collection of *L. plantarum* strains isolated from fermented olives and other plant food types. *L. plantarum* is typically highly abundant in olive fermentations (Hurtado *et al*., 2012). Assessments of the population sizes of individual *L. plantarum* strains in olive fermentations over time have shown how these fermentations are highly dynamic, likely undergoing succession processes at both the species and strain levels (Zaragoza *et al*., 2017). These findings are notable because although LAB have received considerable attention for their contributions to plant fermentations, the diversity, abundance and importance of *L. plantarum* and other LAB in plant microbiomes are not well understood (Yu *et al*., 2020). It has been found that LAB in spontaneous (wild) plant food fermentations are subject to dispersal and selection constraints (Miller *et al*., 2019). However, adaptations expressed by these bacteria that are specific to plant environments and interactions between the same or highly-related LAB species remain to be determined.

*L. plantarum* was isolated from olive fermentations (AJ11, BGM55, BGM37, BGM40, and EL11), tomato fermentations (T2.5 and WS1.1), teff injera fermentations (W1.1, B1.1, and B1.3), wheat sourdough starter (K4), wheat boza (8.1), and prickly pear cactus fruit (1B1) (**Table 1**). Some isolates were collected from the same source either at the same time (strain B1.1 and B1.3) or on different days over the course of fermentation (strains AJ11, BGM37, and BGM40). The strains were selected without considering special criteria or selective pressure. A reference strain from saliva (NCIMB8826R) was used for comparison. To investigate their phenotypic range, the *L. plantarum* strains were evaluated for growth on a variety of plant-associated carbohydrates and during exposure to high NaCl (4% (v/v)), ethanol (EtOH) (8% and 12% (v/v)), or surfactant (sodium dodecyl sulfate (SDS, 0.03% (w/v)) stress. The isolates were measured for the capacity to grow at a low pH (pH 3.5) as well as survive (pH 2) and tolerate a high temperature (50 °C) incubation. Biofilm formation and growth inhibition of *Saccharomyces cerevisiae* UCDFST 09-448, a pectinolytic spoilage yeast (Golomb *et al*., 2013), were also tested. Lastly, to establish the genetic basis for the observed strain differences, multi-locus sequence typing (MLST) and comparative genomics were performed.

**Table 1.** *L. plantarum* strains used in this study.

| **Strain name** | **Isolation source** | **Isolation date <sup>a</sup>** | **References** |
| --- | --- | --- | --- |
| **AJ11** | Fermented olives;<br>commercial fermentation | 12/02/2010 | Golomb *et al.* 2013 |
| **BGM55** | Fermented olives; pilot-<br>scale fermentation<br>inoculated with *S.*<br>*cerevisiae* 09-448 | 03/07/2011 | Golomb *et al.* 2013 |
| **BGM37** | Olive fermentation brine;<br>commercial fermentation | 01/04/2011 | Golomb *et al.* 2013 |
| **BGM40** | Fermented olives;<br>commercial fermentation | 01/26/2011 | Golomb *et al.* 2013 |
| **EL11** | Fermented olives;<br>commercial fermentation | 12/04/2009 | Golomb *et al.* 2013 |
| **K4** | Wheat sourdough starter | 09/15/2014 | This study |
| **8.1** | Wheat boza | 09/15/2014 | This study |
| **W1.1** | White flour teff injera | 04/04/2015 | This study |
| **B1.1** | Brown flour teff injera | 04/04/2015 | This study |
| **B1.3** | Brown flour teff injera | 04/04/2015 | This study |
| **T2.5** | Fermented tomatoes | 08/20/2015 | This study |
| **WS1.1** | Fermented tomatoes<br>(spoiled) | 08/20/2015 | This study |
| **1B1** | Ripe cactus fruit (*Opuntia*<br>*ficus-indicia*) | 10/25/2011 | Tyler *et al.* 2016 |
| **NCIMB8826R** | Human saliva | N/A | Yin *et al.* 2018 |

<sup>a</sup> Month/Day/Year. N/A, not available

## Results

### Strain differentiation and phylogenetic analysis

The isolates were identified as *L. plantarum* by 16S rRNA gene sequence analysis and differentiated from the closely-related species *Lactiplantibacillus pentosus* (formerly *Lactobacillus pentosus* (Zheng *et al*., 2020)) and *Lactiplantibacillus paraplantarum* (formerly *Lactobacillus paraplantarum* (Zheng *et al*., 2020)) by multiplex PCR targeting *recA* (Torriani *et al*., 2001).

The strains were also found to have unique allelic MLST sequence types (ST) (**Table S1**), thus confirming that they are genetically distinct and not derived from the same clonal populations. Among the eight genes tested by MLST, between 6 (*uvrC*) and 12 (*pyrG*) different alleles were found (**Table S1**). Phylogenetic analysis of the ST showed that the *L. plantarum* strains clustered into two clades (**Fig. 1A**). The isolates from fermented olives were contained in one clade, suggesting they are more closely related to each other and to the teff injera strain B1.3 than those retrieved from other sources. The two other strains from teff injera (B1.1 and W1.1) clustered together in the other clade which also contained NCIMB8826R and the strains isolated from wheat boza, sourdough, cactus fruit, and fermented tomatoes (**Fig. 1A**). When examined in a MLST phylogenetic tree containing 264 other *L. plantarum* strains (**Fig. S1**), the *L. plantarum* isolates collected from fermented olives remained clustered closely together, whereas the others were distributed across the tree.

**Fig. 1.**
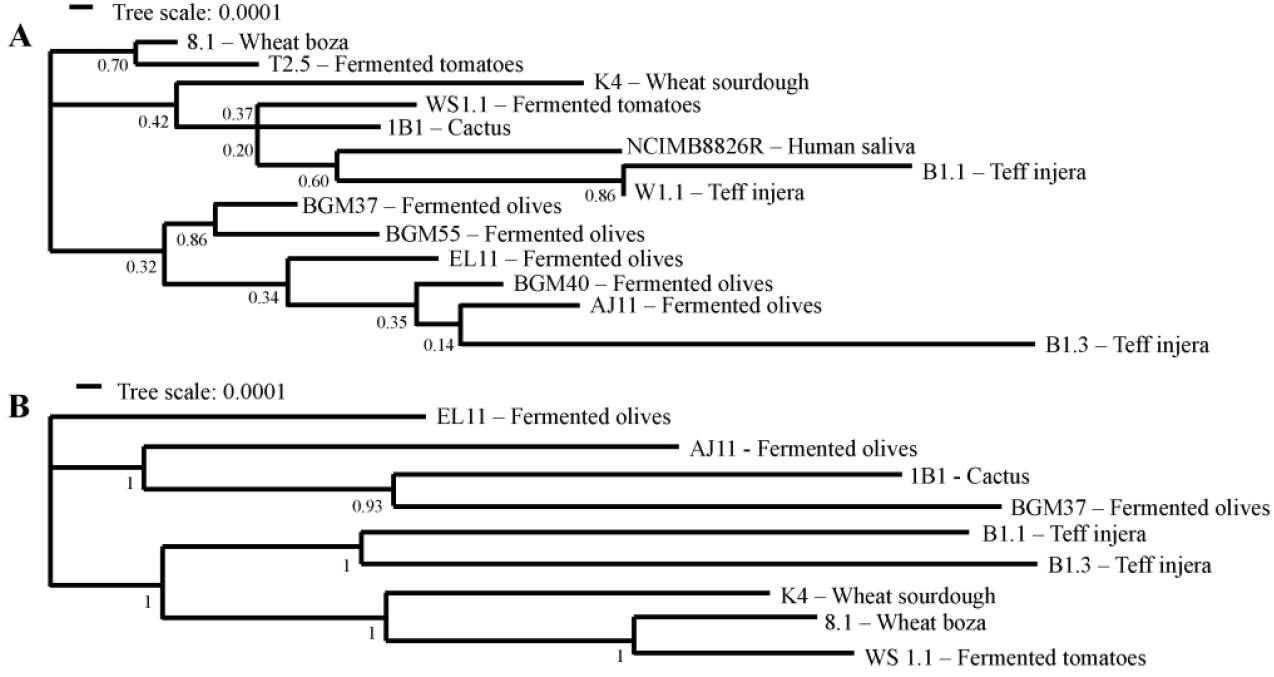
Phylogenetic relationships between *L. plantarum* strains. **(A)** Phylogenetic relationships of 14 strains of *L. plantarum* based on MLST profiles with *pheS, pyrG, uvrC, recA, clpX, murC, groEL*, and *murE* (**Table S8**) and **(B)** nine *L. plantarum* strains based on concatenated core protein sequences using the maximum likelihood method with bootstrap values calculated from 500 replicates using MEGA (7.0) (Kumar et al., 2016).

### Carbohydrate utilization capacities

The capacity of the *L. plantarum* strains to use different sugars for growth was measured using MRS, a complete medium commonly used for cultivation of LAB (De Man et al., 1960). To exclude metabolizable carbon sources, the MRS was modified (mMRS) to remove beef extract and dextrose. In mMRS containing glucose, maltose, or sucrose, all *L. plantarum* strains except B1.3 (teff injera) and 8.1 (wheat boza) were found to have robust growth according to area under the curve (AUC) rankings (**Fig. 2, Fig. 3, and Table S2**). Those strains which grew robustly reached maximum OD_600_ values within 12 h (**Fig. 3 and Table S3**) and displayed growth rates ranging from a low of 0.31 ± 0.01 h^-1^ (strain W1.1 (teff injera) in maltose) to a high of 0.45 ± 0.01 h^-1^ (BGM37 (fermented olives) in glucose) (**Table S4**). By comparison, the growth rate of B1.3 was lower in glucose (0.20 ± 0.00 h^-1^) and maltose (0.15 ± 0.00 h^-1^) compared to the other strains (**Fig. 3 and Table S4**). In mMRS-sucrose, both B1.3 and 8.1 exhibited poor growth (**Fig. 2, Fig. 3, and Table S2**).

**Fig. 2.**
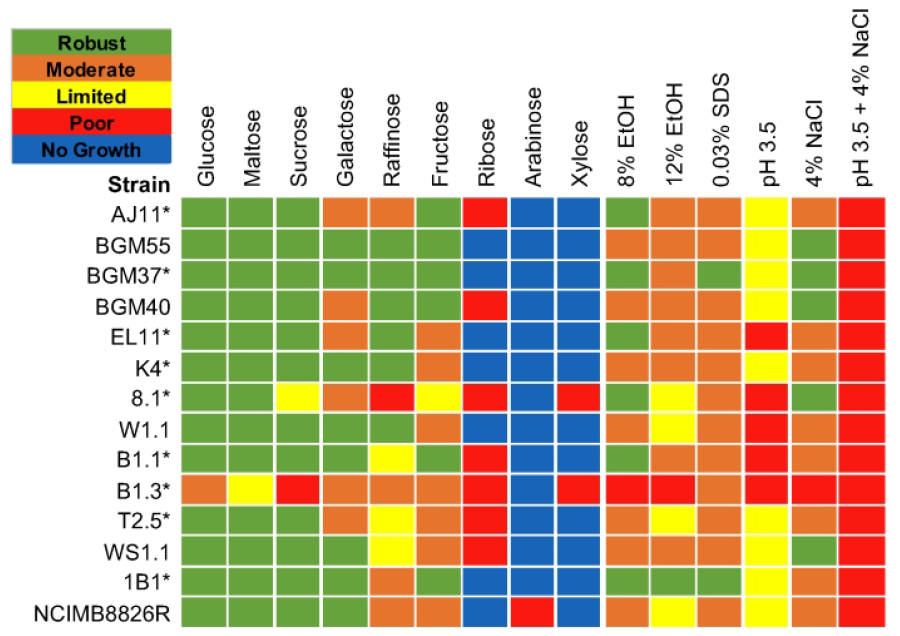
*L. plantarum* phenotype profiles. Area under the curve (AUC) values were used to illustrate *L. plantarum* capacities to grow in mMRS containing different sugars and in mMRS-glucose in the presence of 8% (v/v) EtOH, 8% (v/v) EtOH and then 12% (v/v) EtOH (12% EtOH), 0.03% (w/v) SDS, 4% (w/v) NaCl or set at pH 3.5 without or with 4% (w/v) NaCl. AUC values for the growth curves were ranked as “robust” (AUC between 150 and 115), “moderate” (AUC between 114 and 80), “limited” (AUC between 79 and 45), “poor” (AUC < 45), or “no growth” (AUC was equivalent to the strain growth in mMRS lacking a carbohydrate source). *L. plantarum* growth in mMRS-glucose supplemented with an equal volume of water instead of EtOH, NaCl, or SDS was not significantly different compared to growth in mMRS-glucose (p > 0.05). * indicates strains examined by whole genome sequencing.

**Fig. 3.**
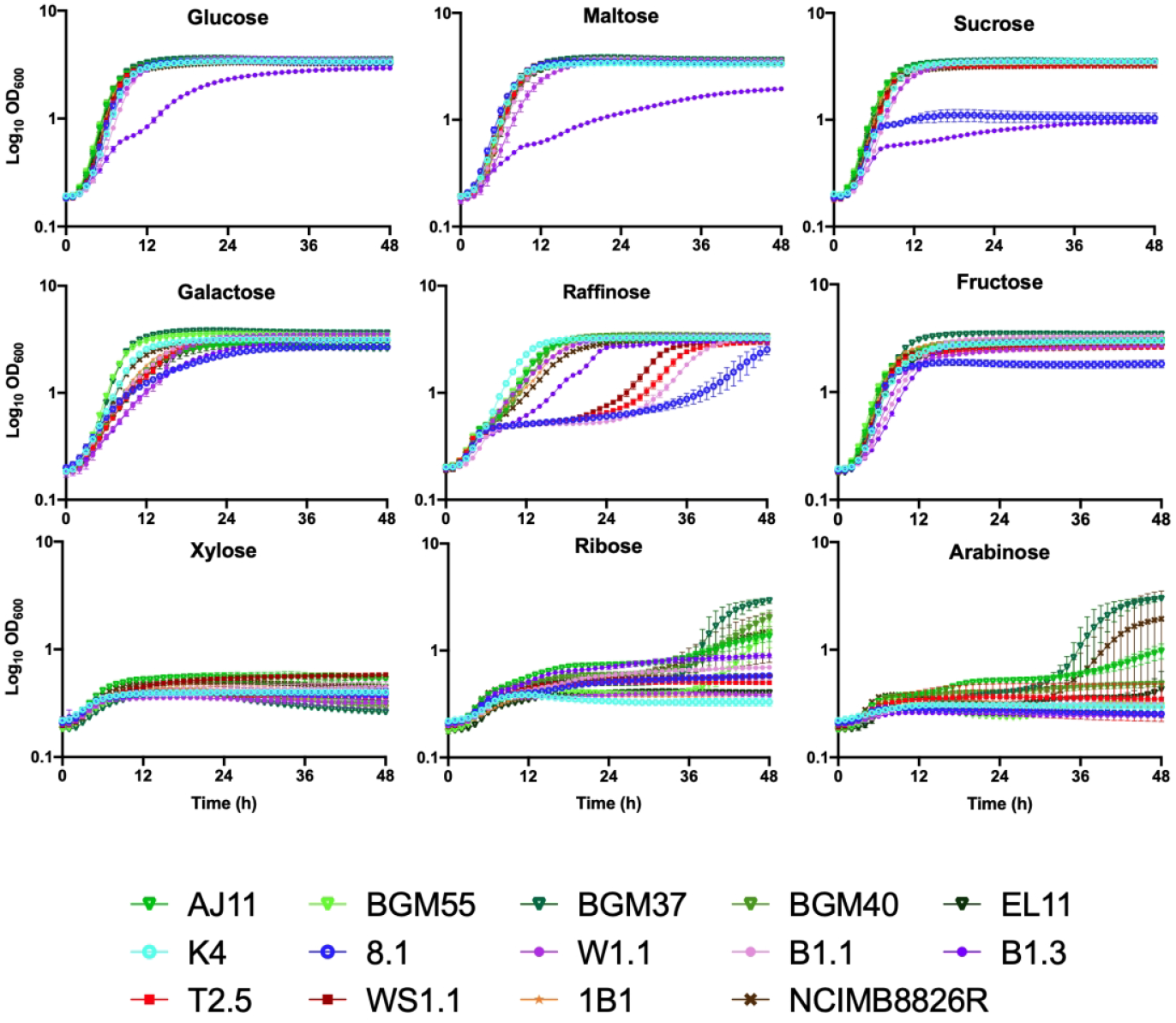
Growth of *L. plantarum* in mMRS containing different mono-, di-, and tri-saccharides. *L. plantarum* was incubated in mMRS containing 2% (w/v) of each sugar at 30 °C for 48 h. The avg ± stdev OD600 of three replicates for each strain are shown.

All strains grew moderately to robustly when galactose was provided as the sole carbon source in mMRS (**Fig. 2, Fig. 3, and Table S2**). Growth rates ranged from a low of 0.16 ± 0.003 h^-1^ (B1.3 (teff injera)) to a high of 0.42 ± 0.01 h^-1^ (BGM37 (fermented olives)) (**Table S4**). Final OD_600_ values measured after 24 h incubation ranged from 2.58 ± 0.05 (BGM40 (fermented olives)) to 3.62 ± 0.03 (BGM37) (**Table S3**). Because incubation in glucose-containing MRS prior to exposure to mMRS-galactose might result in carbon catabolite repression (Kremling *et al*., 2015), several strains with only moderate growth in that culture medium (AJ11, BGM40, and EL11 (fermented olives), 8.1 (wheat boza), B1.3 (teff injera), and T2.5 (fermented tomatoes)) were inoculated in succession into mMRS-galactose. However, prior exposure to mMRS-galactose did not result in higher AUC values (data not shown).

In mMRS with raffinose, all five *L. plantarum* strains isolated from fermented olives (BGM37, BGM55, BGM40, AJ11, and EL11) exhibited either moderate or robust growth (**Fig. 2, Fig. 3, and Table S2, S3, and S4**). Although strain W1.1 (teff injera) also grew robustly, the other strains isolated from teff and wheat fermentations (8.1, B1.1, and B1.3) and both strains isolated from fermented tomatoes (T2.5 and WS1.1) displayed limited or poor growth (**Fig. 2, Fig. 3 and Table S2, S3, and S4**). To address whether the poor growth of those isolates was due to carbon-catabolite repression, serial passage in mMRS-raffinose was performed. Notably, growth of four out of the five strains (B1.1, 8.1, T2.5, and WS1.1) was improved by successive cultivation in mMRS-raffinose (**Fig. S2**).

When fructose was provided, all *L. plantarum* isolates except for strain 8.1 (wheat boza) exhibited either moderate or robust growth (**Fig. 2, Fig. 3, and Table S2**). Similar to incubation in glucose and galactose, strain BGM37 (fermented olives) reached the highest OD_600_ (OD_600_ = 3.44 ± 0.03) (**Table S3**). Remarkably, growth of B1.3 (teff injera) was improved in mMRS-fructose compared to the other sugars tested, as demonstrated by a higher growth rate (**Table S4**) and final OD_600_ (**Table S3**). Similar to the lack of effect on AUC values found after successive passage in the presence of mMRS-galactose, no significant differences in growth were found for any of the 14 strains after multiple passages in mMRS-fructose (data not shown).

Growth of *L. plantarum* was poor in mMRS containing xylose, ribose, or arabinose. Only four olive-associated strains (AJ11, BGM55, BGM37, and BGM40) and NCIMB8826R grew in the presence of mMRS-ribose or mMRS-arabinose and none grew in mMRS-xylose (**Fig. 2, Fig. 3, and Table S2, S3, and S4**). After 38 h in mMRS-ribose, the OD_600_ values for those strains ranged from a low of 1.36 ± 0.15 (AJ11 (fermented olives)) to a high of 2.93 ± 0.17 (BGM37 (fermented olives)) (**Fig. 3 and Table S3**). In mMRS with arabinose, only NCIMB8826R and BGM37 grew, reaching an OD_600_ of 1.94 ± 1.56 and 2.99 ± 0.14, respectively (**Fig. 3 and Table S3**). To investigate whether growth could be improved by prior exposure to those pentose sugars, strains AJ11, BGM37, 8.1, and NCIMB8826R were incubated with successive passages in mMRS-ribose or mMRS-arabinose. This resulted in shorter lag phase times and higher final OD_600_ values for AJ11, BGM37, and NCIMB8826R in both media (**Fig. S3 and Fig. S4**). By comparison, no difference in growth was observed for strain 8.1 (wheat boza) in mMRS-ribose or mMRS-arabinose irrespective of the adaptation period (**Fig. S3 and Fig. S4**).

### Growth in the presence of EtOH

Because mMRS-glucose resulted in robust growth of the majority of *L. plantarum* strains investigated here, that culture medium was used for investigation of stress tolerance properties. In mMRS-glucose containing 8% (v/v) (174 mM) ethanol (EtOH), the AUCs for all strains except B1.3 (teff injera) were either moderate or robust (**Fig. 2, Fig. 4, and Table S2**). Although lag phase times were longer (data not shown) and growth rates were reduced when EtOH was included in the culture medium (**Table S5**), the growth curves of six strains (AJ11, BGM37, and EL11 (fermented olives), 8.1 (wheat boza), B1.1 (teff injera), and 1B1 (cactus fruit)) were still regarded as robust according to AUC assessments (**Fig. 2**). Surprisingly, two strains, BGM37 (fermented olives) and 1B1 (cactus fruit), reached a higher final OD_600_ in mMRS-glucose with 8% (v/v) EtOH than in mMRS-glucose alone (Student t-test, P < 0.05) (**Table S3**).

**Fig 4.**
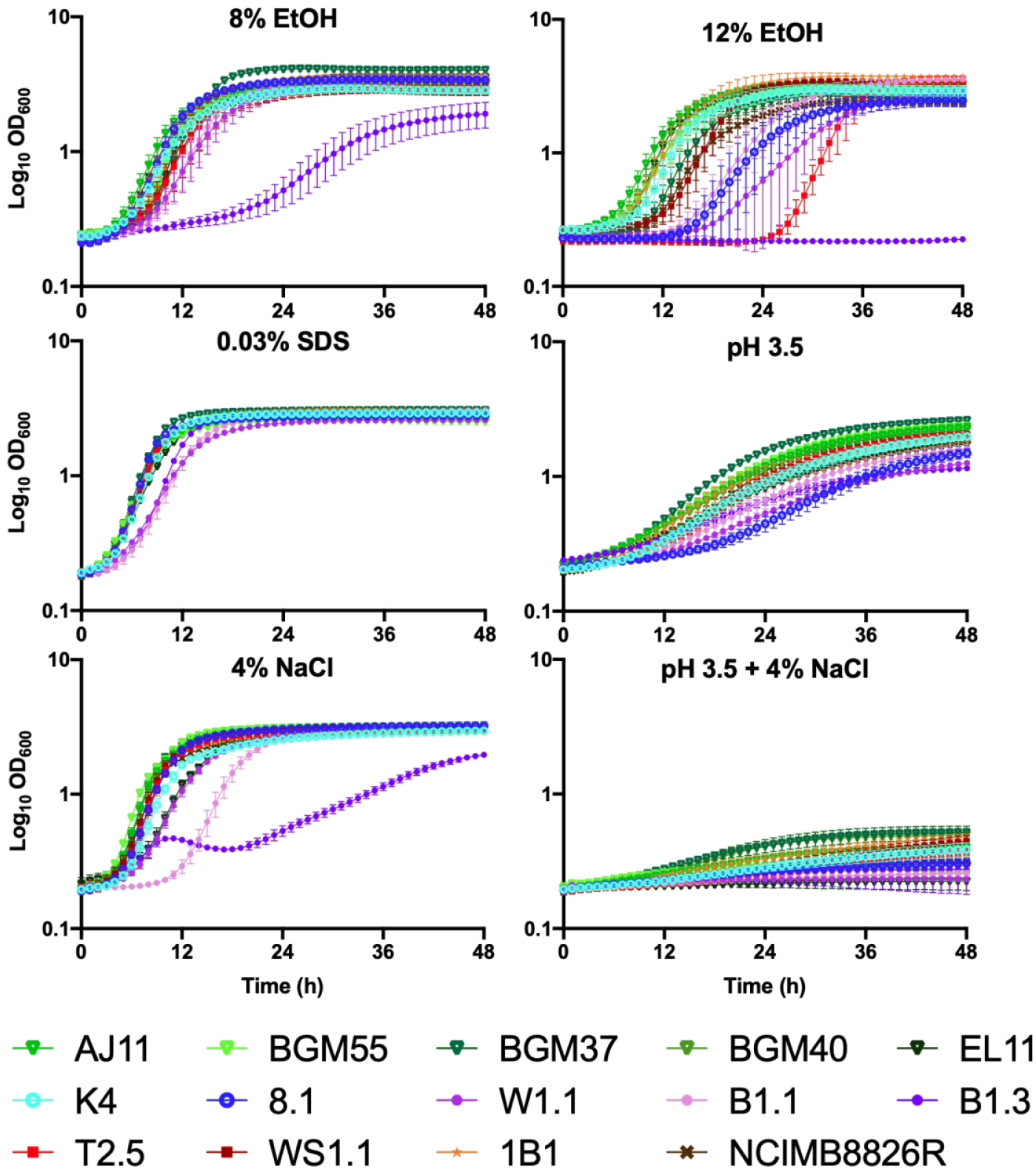
Growth of *L. plantarum* in mMRS-glucose exposed to different environmental stressors. *L. plantarum* was incubated in mMRS-glucose containing 8% (v/v) EtOH, 12% (v/v) EtOH, 0.03% (w/v) SDS, or 4% (w/v) NaCl with or without adjustment to pH 3.5 and incubated at 30 °C for 48 h. The avg ± stdev OD600 of three replicates for each strain are shown.

None of the *L. plantarum* strains tested here were able to grow over a 48 h period when incubated directly in mMRS-glucose with 12% (v/v) (260 mM) EtOH (data not shown). To determine whether a more gradual exposure to high EtOH concentrations would change this outcome, the strains were incubated in mMRS-glucose containing 8% (v/v) EtOH overnight prior to inoculation into mMRS-glucose with 12% (v/v) EtOH. This modification resulted in robust growth of 1B1 (cactus fruit) (**Fig. 2, Fig. 4, and Tables S2, S3, and S5**). Eight other strains (AJ11, BGM55, BGM37, BGM40, and EL11 (fermented olives), K4 (wheat sourdough), B1.1 (teff injera), and WS1.1 (fermented tomatoes)) exhibited moderate growth according to AUC values as a result of the step-wise transfer to the higher (12% (v/v)) EtOH conditions (**Fig. 2, Fig. 4, and Tables S2, S3, and S5**).

### Growth in the presence of detergent (SDS) stress

While most of the *L. plantarum* strains exhibited moderate growth when SDS (0.03% (w/v) (0.10 mM)) was included in mMRS-glucose, two strains BGM37 (fermented olives) and 1B1 (cactus fruit) grew robustly (**Fig. 2, Fig. 4, and Tables S2, S3, and S5**). Remarkably, the growth rate of strain B1.3 (teff injera) was higher in the presence of SDS (0.32 ± 0.003 h^-1^) (**Table S5**) as opposed to its absence (0.20 ±0.003 h^-1^) (**Table S4**) and it reached a higher AUC (107 ± 0.17) (**Table S2**).

### Growth at pH 3.5 and in the presence of 4% NaCl

Growth of *L. plantarum* was reduced in mMRS-glucose adjusted to a pH of 3.5 (**Fig. 2, Fig. 4, and Table S2)**. However, the strains isolated from brine-based, fruit fermentations (AJ11, BGM55, BGM37, BGM40, and EL11 (fermented olives) and T2.5 and WS1.1 (fermented tomatoes)), grew significantly better under those conditions compared to the *L. plantarum* isolated from grain fermentations (Student T-test, p < 0.05). The strains from grain-based fermentations (K4 (wheat sourdough), 8.1 (wheat boza), W1.1, B1.1, and B1.3 (teff injera)) grew poorly in the acidified mMRS (pH 3.5) (**Fig. 2, Fig. 4, and Table S2**), yielding low growth rates (0.06 ± 0.01 h^-1^) (**Table S5**) and final OD_600_ values (1.50 ± 0.19) (**Table S3**).

When 4% (w/v) NaCl was included in mMRS-glucose, five strains isolated from different sources (BGM55, BGM37, and BGM40 (fermented olives), 8.1 (wheat sourdough), and WS1.1 (fermented tomatoes)) were classified as robust according to their AUC values (**Fig. 2** and **Table S2**). The growth of strain B1.3 (teff injera) was the most negatively impacted by the addition of salt into the laboratory culture medium (**Fig. 4 and Tables S2, S3, and S5**).

All *L. plantarum* strains were inhibited in mMRS-glucose containing 4% (w/v) NaCl and a starting pH of pH 3.5 (**Fig. 2, Fig. 4, and Table S2**). The final OD_600_ values ranged from a low of 0.23 ± 0.00 (W1.1, teff injera) to a high of 0.52 ± 0.06 (BGM37, fermented olives) (**Table S3**). Although the AUCs of all strains were regarded to be poor, growth rates of those isolated from brine-based, fruit fermentations (AJ11, BGM55, BGM37, BGM40, and EL11 (fermented olives) and T2.5 and WS1.1 (fermented tomatoes)) were significantly higher than those isolated from grain-based fermentations (K4 (wheat sourdough), 8.1 (wheat boza), W1.1, B1.1, and B1.3 (teff injera)) (p < 0.05, Student’s T-test).

### Survival at pH 2

Within 15 min incubation in physiological saline adjusted to pH 2, a 10^4^ to 10^6^ -fold reduction in cell viability was observed (**Fig. 5A**). After 30 min exposure to pH 2, strains B1.3 (teff injera), BGM40 (fermented olives), and NCIMB8826R (saliva, reference strain) were no longer detectable by colony enumeration. BGM37 (fermented olives), B1.1 (teff injera), and T2.5 (fermented tomatoes) were no longer viable by 60 min (**Fig. 5A**). *L. plantarum* AJ11, BGM55, and EL11 (fermented olives), 8.1 (wheat boza), and WS1.1 (fermented tomatoes) exhibited the highest acid tolerance and were still viable according to colony enumerations performed on cells collected after 60 min incubation. Unlike the findings for growth under acidic conditions (pH 3.5) (**Fig. 4**), there were no obvious isolation-source dependent trends in *L. plantarum* strain survival.

**Fig 5.**
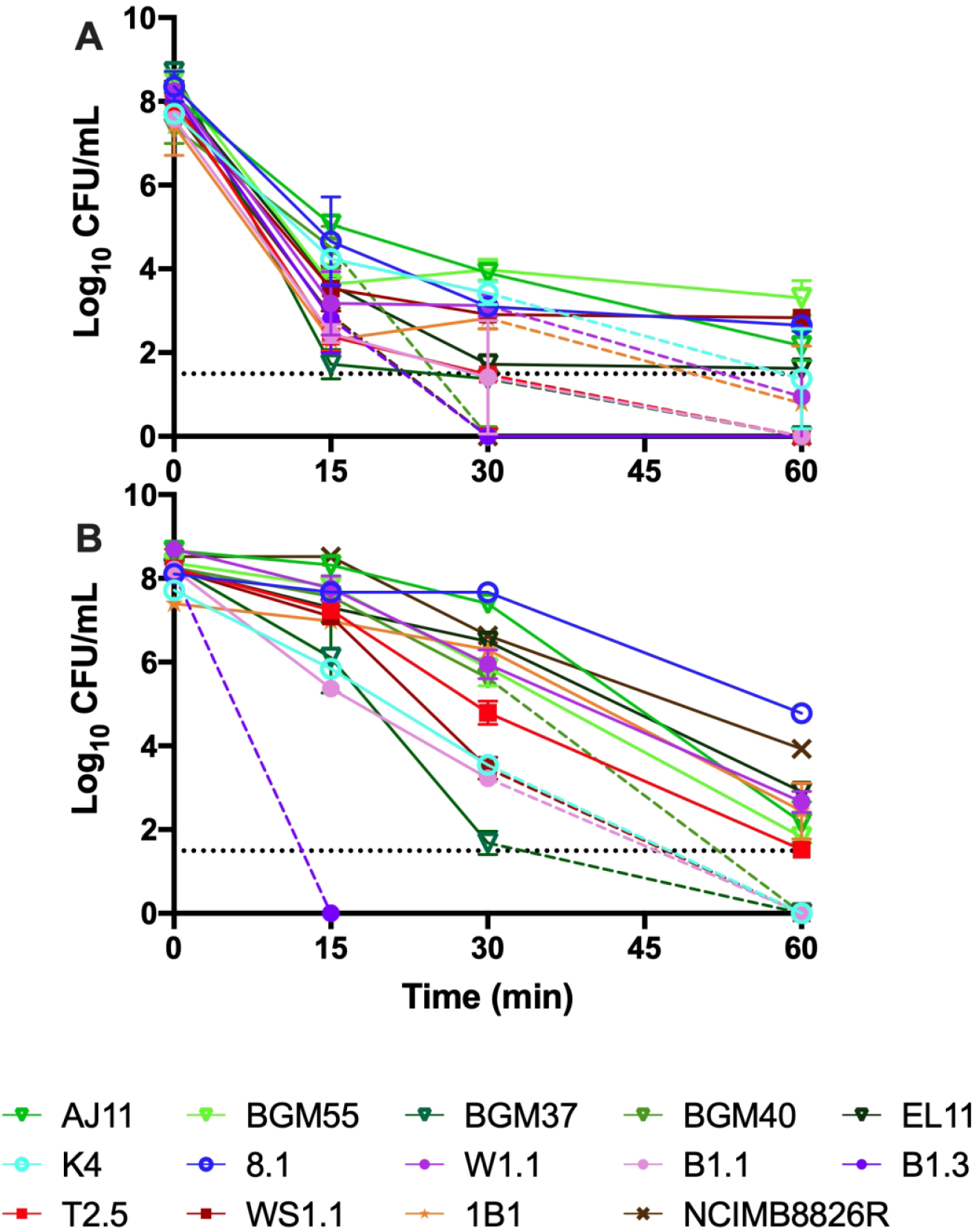
Survival of *L. plantarum* at (A) pH 2 and at (B) 50 °C. **(A)** Viable cells were enumerated after 0, 15, 30, and 60 min of incubation in physiological saline at pH 2 or **(B)** in PBS at 50 °C. The dashed lines indicate when the number of viable cells were below the detection limit (34 CFU/mL). The avg ± stdev CFU/mL values of three replicates for each strain are shown.

### Survival at 50 °C

Survival at 50 °C spanned a 10^6^ - fold range (**Fig. 5B**). Viable B1.3 (teff injera) cells were no longer detected after incubation at 50 °C for 15 min (1 × 10^8^ cells/ml present in the inoculum). After 60 min, AJ11 and EL11 (fermented olives), 8.1 (wheat boza), W1.1 (teff injera), 1B1 (cactus fruit), and NCIMB8826R (saliva, reference strain) were still culturable in a range from 5 × 10^4^ (8.1) to 1.5 × 10^2^ (AJ11) CFU/ml, spanning a 10^3^ - to 10^6^ -fold reduction in viable cell numbers (**Fig. 5B**). Similar to survival to pH 2, no obvious isolation-source dependent differences in survival were observed.

### Biofilm forming capacity

Because biofilm formation is an indicator of bacterial capacities to tolerate environmental stress (Yin *et al*., 2019) and *L. plantarum* biofilm formation is partially dependent on carbon source availability (Fernández Ramírez *et al*., 2015), we examined the capacity of *L. plantarum* to produce biofilms during growth in mMRS with glucose, fructose, or sucrose. Only BGM55 and BGM37 (fermented olives), 8.1 (wheat boza), W1.1 and B1.1 (teff injera), T2.5 and WS1.1 (fermented tomatoes) formed robust biofilms after growth in at least one of those laboratory culture media **(Fig. 6**). Whereas injera strain W1.1 only developed a biofilm when grown in mMRS-fructose, the other isolates formed robust biofilms in the presence of at least two different sugars (**Fig. 6**). Both strains isolated from fermented tomatoes, T2.5 and WS1.1, formed extensive biofilms when grown in the presence of either glucose or fructose. Notably, biofilm formation was not associated with robust strain growth. Strain 8.1 formed a biofilm in mMRS-sucrose (**Fig. 6**) despite showing poor growth (**Fig. 2**) and reaching a low final OD_600_ (**Table S3**) in that culture medium. Conversely, strain K4 grew well in mMRS-sucrose but did not produce a biofilm.

**Fig 6.**
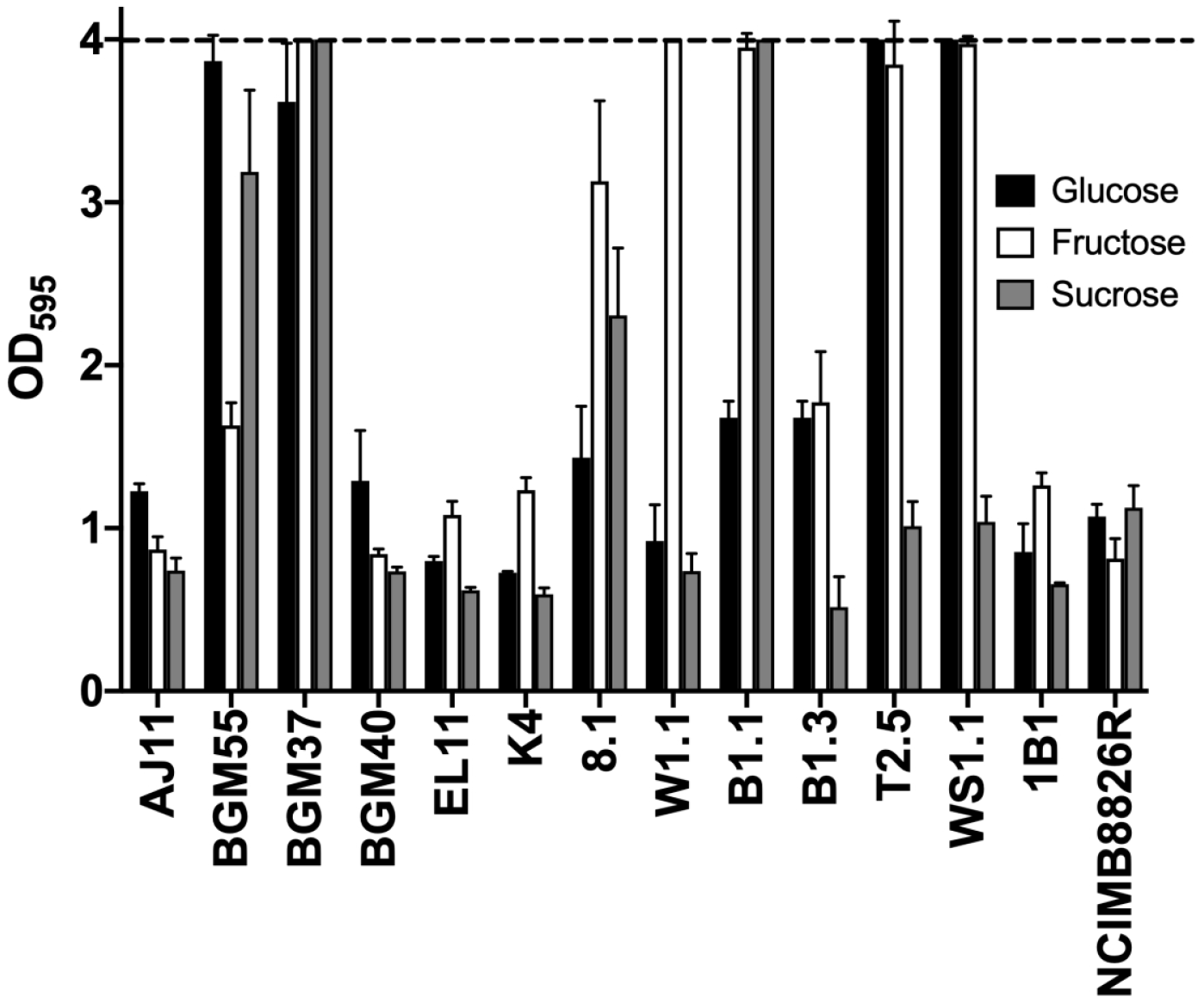
*L. plantarum* biofilm formation during growth in mMRS with glucose, fructose, or sucrose. *L. plantarum* was incubated in mMRS-glucose, mMRS-fructose, and mMRS-sucrose in 96-well, polystyrene microtiter plates at 30 °C for 48 h. The non-adherent cells were removed by washing with PBS. The remaining cells were stained with 0.05% crystal violet (CV). OD595 values of wells without cells did not exceed 0.22. The upper detection limit as indicated by the stippled line was an OD595 of 4.0. The avg ± stdev OD595 of three replicate wells after CV staining are shown.

### Antifungal activity of *L. plantarum* cell-free culture supernatant (CFCS)

Growth rates and final OD_600_ values of *S. cerevisiae* UCDFST 09-448 were reduced when incubated in the presence of the *L. plantarum* CFCS (**Table S6)**. All *L. plantarum* CFCSs inhibited *S. cerevisiae* growth, however there were some strain-specific differences (**Fig. 7 and Table S6**). Collectively the CFCSs from strains isolated from fermented olives (AJ11, BGM55, BGM37, BGM40, EL11) and fermented tomatoes (WS1.1 and T2.5) were significantly (p < 0.05, Student’s T-test) more inhibitory than those isolated from fermented grains (K4, 8.1, W1.1, B1.1, and B1.3). Growth inhibition resulting from exposure to the CFCS from olive strains ranged between 29.8% ± 4.87 (BGM55) to 34.1% ± 9.4 (BGM40). By comparison, growth inhibition with CFCS from *L. plantarum* isolated from grain fermentations was only between 20.1% ± 1.06 (B1.1) to 22.68% ± 1.46 (8.1). Interestingly, the growth pattern of *S. cerevisiae* in the presence of teff injera strain B1.3 CFCS (31.4 ± 1.27) was more similar to strains from fermented olives than grains.

**Fig 7.**
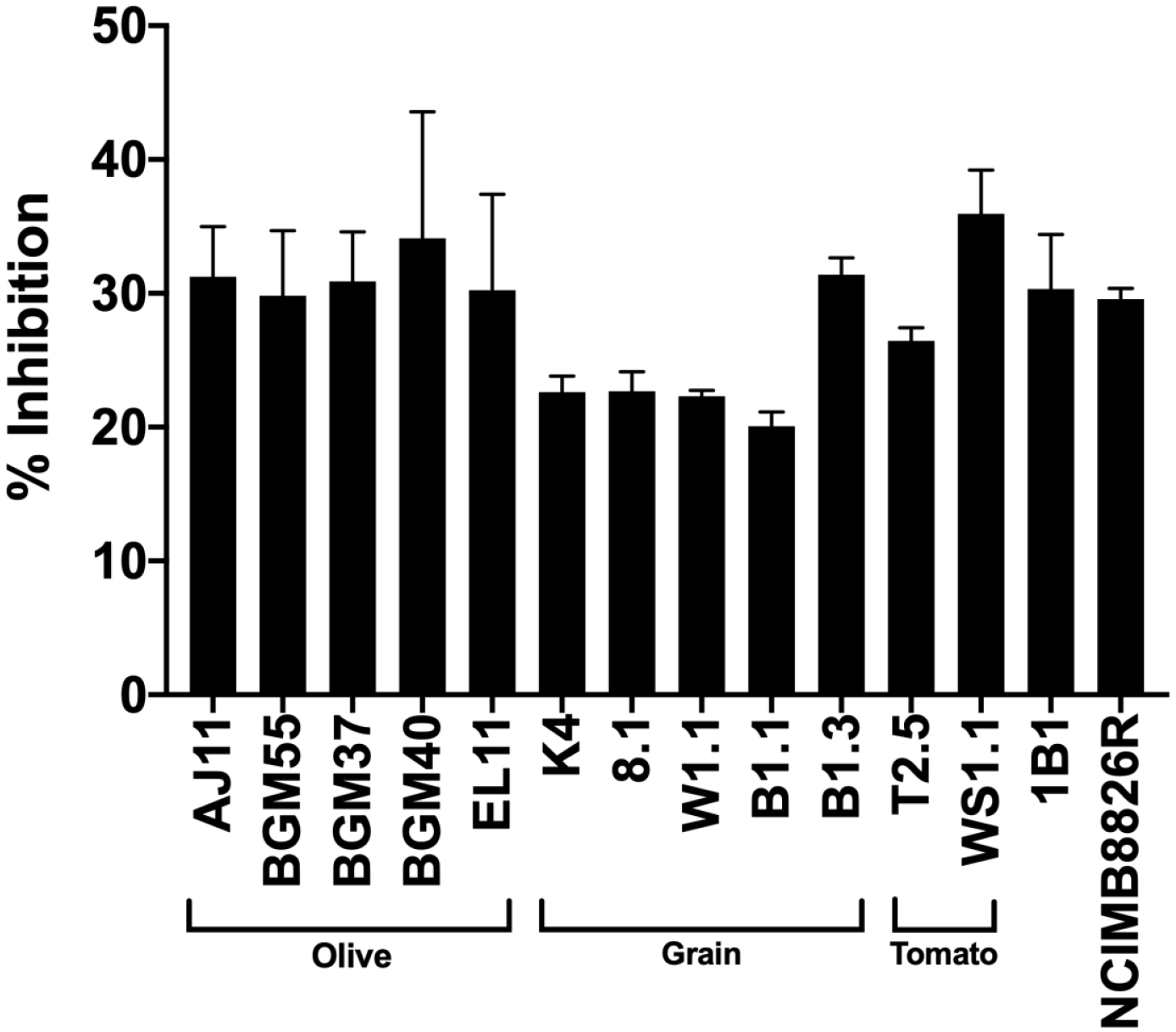
*S. cerevisiae* growth inhibition in the presence of *L. plantarum* CFCS. *S. cerevisiae* UCDFST- 09-448 was incubated in a 1:1 ratio of 2X YM and pH adjusted (pH 3.8) *L. plantarum* CFCS from cMRS. Growth was measured by monitoring the change in OD_600_ over 24 h. Percent inhibition was determined by comparing the final OD600 of *S. cerevisiae* grown in the presence of CFCS to growth in a 1:1 ratio of 2X YM and pH adjusted (pH 3.8) cMRS.

### Comparisons of *L. plantarum* genomes

Nine of the fourteen strains were selected for genome sequencing (PacBio or Illumina platforms) based on the variations in their phenotypic profiles (**Fig. 2**). Genome assembly for strains sequenced using PacBio resulted in fewer contigs (min of 3 and max of 9) and higher coverage (min of 140X and max of 148X) compared to Illumina (contigs: min of 29 and max of 120; coverage (min of 27X and max of 128X) (**Table 2**). Genome sizes ranged from 3.09 Mbp (B1.3 (teff injera)) to 3.51 Mbp (WS1.1 (fermented tomatoes)) and total numbers of predicted coding sequences ranged from 3,088 (K4 (wheat sourdough)) to 3,613 (WS1.1) (**Table 2**).

**Table 2.** *L. plantarum* genome coverage and assembly statistics.

| **Strain <sup>a</sup>** | **Accession No.** | **Genome Size (Mb)** | **# of Contigs** | **Coverage** | **N50** | **L50** | **% GC Content** | **# of CDS** |
| --- | --- | --- | --- | --- | --- | --- | --- | --- |
| **AJ11** | WWDD000000000 | 3.27 | 29 | 27X | 252487 | 6 | 44.54 | 3,214 |
| **BGM37** | WWDC000000000 | 3.46 | 46 | 66X | 155998 | 7 | 44.15 | 3,467 |
| **EL11** | WWDB000000000 | 3.28 | 29 | 128X | 1944449 | 5 | 44.31 | 3,231 |
| **K4** | WWDF000000000 | 3.16 | 3 | 148X | 3157988 | 1 | 44.60 | 3,088 |
| **8.1** | WWDE000000000 | 3.37 | 9 | 140X | 3066287 | 1 | 44.40 | 3,366 |
| **B1.1** | WWCZ000000000 | 3.17 | 120 | 76X | 59472 | 19 | 44.55 | 3,242 |
| **B1.3** | WWCY000000000 | 3.09 | 5 | 145X | 2939357 | 1 | 44.50 | 3,157 |
| **WS1.1** | WWDA000000000 | 3.51 | 99 | 30X | 78600 | 12 | 44.11 | 3,613 |
| **1B1** | WWDG000000000 | 3.34 | 60 | 28X | 109565 | 11 | 44.34 | 3,371 |

<sup>a</sup> The genomes of AJ11, EL11, BGM37, WS1.1, and 1B1 were sequenced by Illumina MiSeq V2 (2 X 250). The genomes of strains
K4, 8.1, and B1.3 were sequenced by PacBio RSII (P6-C4 sequencing chemistry).

The core- and pan-genomes of the nine strains consisted of 2,222 and 6,277 genes, respectively (**Fig. S5**), numbers consistent with previous comparisons examining larger collections of *L. plantarum* strains (Siezen *et al*., 2010; Martino *et al*., 2016; Choi *et al*., 2018). Alignments of the predicted amino acid sequences for the genes in the core genomes indicated that strains isolated from grain fermentations (K4 (wheat sourdough), 8.1 (wheat boza) and B1.1, and B1.3 (teff injera)) and strain WS1.1 from fermented tomatoes are more closely related to each other than isolates from olives and cactus fruit (**Fig. 1B**). B1.1 and B1.3, two strains originating from the same sample of teff injera, were also shown to share similar core genomes (**Fig. 1B**).

Just as strains 8.1 (wheat boza) and WS1.1 (fermented tomatoes) were found to have similar core genomes (**Fig. 1B**), those two strains are similar according to hierarchical clustering based on the numbers of genes in individual COG categories (**Fig. 8**). The three strains isolated from olives formed a separate clade from those recovered from other sources and were shown to have higher numbers of genes in the carbohydrate metabolism and transport (G) and transcription (K) COGs. *L. plantarum* BGM37, a strain from olives that exhibited the most robust growth on the different carbohydrates compared tested here (**Fig. 2**), also contained the highest numbers of gene clusters annotated to the carbohydrate metabolism and transport COG (256 gene clusters, **Fig 8. and Table S7**).

**Fig 8.**
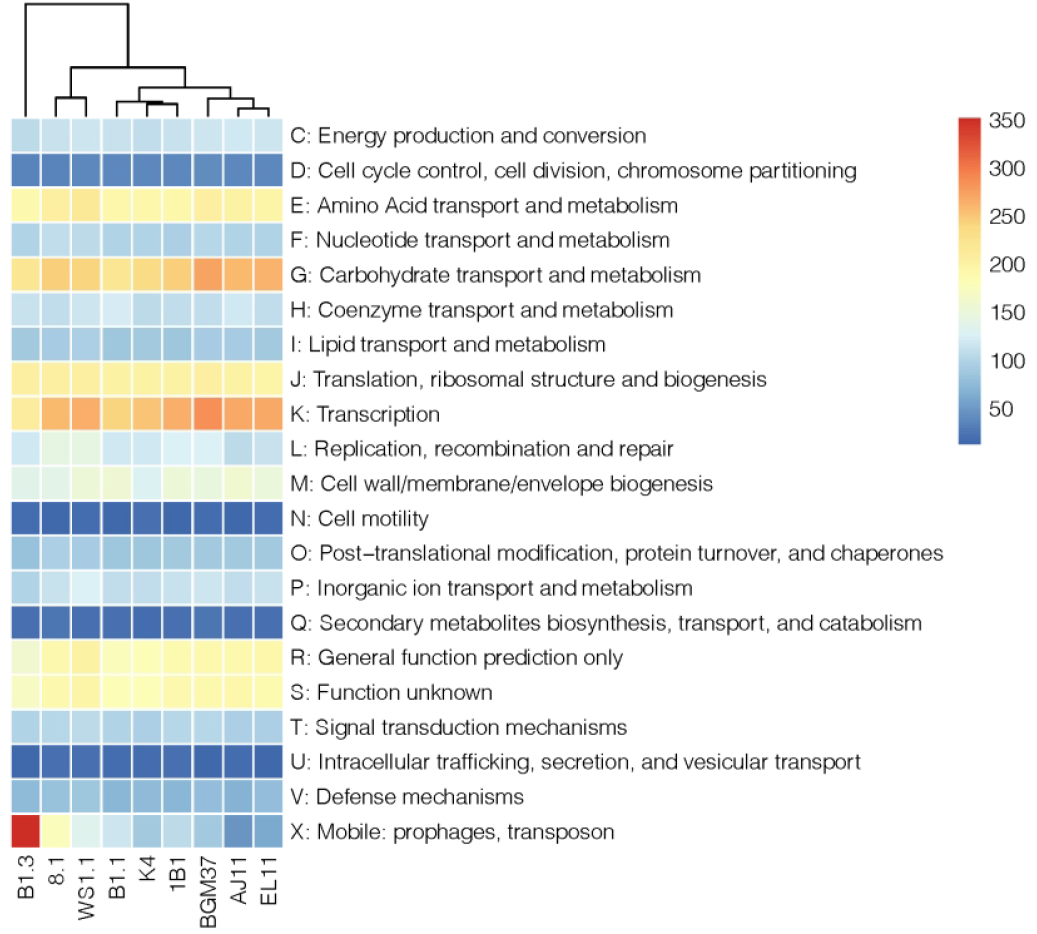
Distribution of COG Categories across *L. plantarum* genomes. Hierarchical clustering of *L. plantarum* based on the number of gene clusters assigned to each functional COG category. Number of gene clusters present in each strain was denoted by the color gradient.

Strain B1.3 was found to contain the lowest number of gene clusters in the carbohydrate metabolism and transport COG (206 gene clusters, **Fig. 8 and Table S7**) and is specifically lacking in several genes required for sugar metabolism and sugar-importing phosphotransferase (PTS) systems (data not shown). Conversely, the genome of B1.3 harbors at least two-fold higher numbers of genes and genetic elements in the mobilome (X) COG compared to the other strains examined (352 gene clusters, **Table S7**). These genomic features include prophages, insertion sequence elements, and transposases that are interspersed throughout the genome and frequently located between genes with known function. For example, a transposon (3.8 kb) is located between the glucose-6-phosphate isomerase (lp_2502) and glucose/ribose porter family sugar transporter (lp_2503) genes that are annotated to be associated with glucose metabolism. Other genes were not present in the B1.3 genome such as the sucrose-associated PTS (lp_3819; *pts24BCA*), possibly indicating why this strain exhibited poor growth in mMRS-sucrose. Strain 8.1, the only other *L. plantarum* strain tested here that grew to a limited extent on sucrose (**Fig. 2**), lacks the first 650 bp of *pts1BCA* (lp_0185), a gene in the sucrose phosphoenolpyruvate (PEP)-dependent phosphotransferase system (PTS) (Saulnier *et al*., 2007; Yin *et al*., 2018).

The number of gene clusters in the other COG categories was largely conserved between strains (**Fig. 8 and Table S7**). These COG categories encode pathways required energy metabolism (glycolysis), synthesis of macromolecules (proteins, nucleotides, and lipids), and stress response. The genomes of all nine strains contain genes encoding chaperones (DnaJK, GroEL, GroES, GrpE, ClpB, ClpL), proteases (ClpX, ClpP, ClpE), DNA repair proteins (RecA, UvrABC), and transcriptional regulators (HrcA, CtsR) critical for *L. plantarum* tolerance to numerous environmental stresses (Papadimitriou *et al*., 2016). Although genes required for citrate metabolism (*citCDEF*) were previously found to be associated with EtOH tolerance (Veen *et al*., 2011) and that locus was flanked by mobile elements in several of the *L. plantarum* strains examined here, the presence of those mobile elements was not correlated with EtOH sensitivity.

## Discussion

This study investigated the phenotypic and genetic properties of *L. plantarum* strains from (fermented) plant sources. The findings broadly show that strains obtained from the same or similar plant environments tend to be more genetically related and share similar carbohydrate utilization and stress tolerance capacities. However, there were still significant differences between all strains, irrespective of their source, a result which suggests that *L. plantarum* has adapted for growth in specific habitats (e.g., olive fermentations) but that intraspecific variation of this generalist species may afford the opportunity for *L. plantarum* strain coexistence by niche differentiation.

Our use of growth curve AUC rankings and the monitoring of growth rates and final OD_600_ values provided a detailed view of *L. plantarum* carbohydrate utilization capacities. The majority of strains exhibited robust growth on glucose, maltose, sucrose, and galactose, moderate growth on raffinose and fructose, and only limited to no growth on ribose, arabinose, and xylose. The moderate or poor growth observed for a few strains when incubated the presence galactose or fructose, was likely not due to carbon catabolite repression (Görke and Stülke, 2008; Kremling *et al*., 2015), but rather a lack of enzymatic capacity to utilize those sugars. These conserved carbohydrate consumption patterns are consistent with prior reports on *L. plantarum* isolated from plants and other host-associated sources (Westby *et al*., 1993; Saulnier *et al*., 2007; Siezen *et al*., 2010; Filannino *et al*., 2014; Siragusa *et al*., 2014). The strains tested here were also able to grow in the presence of 0.03% (w/v) SDS and were severely impaired when incubated in mMRS at pH 3.5 with 4% (w/v) NaCl or inoculated directly into mMRS with 12% (v/v) EtOH.

Other findings were strain specific and similarly consistent with reported phenotypic (Parente *et al*., 2010; Siezen *et al*., 2010; Guidone *et al*., 2014; Ferrando *et al*., 2015, 2016; Gheziel *et al*., 2019; Prete *et al*., 2020) and genomic variations (Molenaar *et al*., 2005; Siezen *et al*., 2010; Siezen and van Hylckama Vlieg, 2011; Martino *et al*., 2016; Choi *et al*., 2018) observed for the *L. plantarum* species. We found that *L. plantarum* growth was highly variable following the sequential incubation in 8% (v/v) and then 12% (v/v) EtOH. Strain growth rates in mMRS with 8% (v/v) EtOH were correlated with those observed for mMRS containing 0.03% SDS (r = 0.561, p < 0.05), thereby indicating overlapping mechanisms in *L. plantarum* strain tolerance to membrane-disruptive compounds (Seddon *et al*., 2004; Bravo-Ferrada *et al*., 2015; Mukhopadhyay, 2015). High temperature tolerance also differed between the *L. plantarum* isolates, such that incubation at 50 °C for 60 min resulted in over a 10^5^ - fold range in strain survival. Survival at pH 2 followed a similar trend, such that some strains were no longer culturable after 15 min, while other strains still formed colonies after prolonged (60 min) incubation. Notably, only two strains from olive fermentations (AJ11 and EL11) and 8.1 from boza survived well under both high temperature and low pH conditions. Although, the genomes were found contain chaperones and proteases known to be involved in *L. plantarum* heat and acid shock responses (Corcoran *et al*., 2008; Mills *et al*., 2011), the unique proteins or pathways expressed by those strains which confer heightened stress tolerance remain to be determined.

Despite the conserved and variable aspects of *L. plantarum* carbohydrate utilization and environment stress tolerance phenotypes, there were other remarkable trends associated with strain isolation source. For example, the isolates from acidic, brine-containing ferments (olives and tomatoes) were more resistant to acidic pH (pH 3.5) and high NaCl (4% w/v) concentrations than those recovered from grain fermentations (wheat boza, wheat sourdough, and teff injera).Genome comparisons using concatenated core gene amino acids showed that strains isolated from grain fermentations are more related to each other than those from other sources. Genetic conservation between olive fermentation-associated strains was observed by MLST and COG gene numbers.

The strains from fermented olives also showed the greatest capacity to consume raffinose (a tri-saccharide composed of galactose, glucose, and fructose). It is also notable that two of those isolates (BGM37 and BGM55) grew equally well in mMRS-galactose as in mMRS-glucose. These results are consistent with the findings that olives leaves and roots contain both raffinose (2.7 ± 0.1 µmol) and galactose (4.8 ± 0.3 µmol) (Cataldi *et al*., 2000) and that the fruits contain galactose along with higher concentrations of glucose, mannitol, and fructose (Gómez-González *et al*., 2010). All strains from olive fermentations also exhibited at least moderate or robust growth in mMRS in the presence of 8% (v/v) EtOH, and the CFCSs from those strains resulted in greater inhibition of *S. cerevisiae* UCDFST-09-448 compared to the CFCSs from *L. plantarum* isolated from other environments. Because yeast are normal members of olive fermentation microbiota, the inhibitory capacity may indicate the presence of shared mechanisms required to prevent yeast overgrowth.

Several strains also showed unique properties illustrative of the phenotypic range of the *L. plantarum* species. Among those strains was BGM37 isolated from the brine of fermented olives. This strain exhibited the most robust growth on the carbohydrates tested here, showed the highest tolerance to 8% EtOH and 0.03% SDS, and was able to form biofilms in the presence of glucose, fructose, and sucrose. Compared to the other strains for which genome sequences were obtained, BGM37 was found to have the second largest genome size (3.46 Mbp) after WS1.1 (3.51 Mbp), a magnitude comparable to the other *L. plantarum* strains with large (complete) genomes published at NCBI (maximum of 3.70 Mbp as of Jan 2021).

*L. plantarum* 1B1, a strain isolated from ripe cactus fruit, is notable because of its robust growth in the presence of either SDS or EtOH. Although other studies reported growth of *L. plantarum* in the presence of EtOH (Veen *et al*., 2011, Brizuela *et al*., 2019, Duley, 2004), the capacity to grow well at 12% EtOH is an unusual trait even among oenological-associated *L. plantarum* (Succi *et al*., 2017). Thus, the unique properties of this single isolate from a fresh fruit source may indicate the presence of a broader diversity of LAB present in the carposphere (Yu *et al*., 2020).

Lastly, strain B1.3 from teff injera exhibited the most restrictive carbon utilization capacities and the lowest levels of environmental stress tolerance among all isolates tested. B1.3 grew poorly on glucose and most other carbohydrates, whereas the other strains from teff injera B1.1 and W1.1 exhibited robust growth on a variety of sugars. Limitations in the ability of B1.3 to consume different sugars was also shown by the lower numbers of gene clusters in the B1.3 genome that are responsible for carbohydrate transport and metabolism. The overall smaller genome size of this strain (3.09 Mbp) and high numbers of genes in the mobilome COG potentially indicates that this strain is undergoing genome reduction for habitat specialization as found for other LAB (e.g., *Lactobacillus bulgaricus* (yogurt) (van de Guchte *et al*., 2006), *Lactobacillus iners* (vagina) (France *et al*., 2016), *Apilactobacillus apinorum* (honeybee) (Endo *et al*., 2018)). Remarkably, the higher growth rate of B1.3 in mMRS-fructose and in the presence of SDS indicates it may be fructophilic and capable of withstanding the presence of membrane disrupting compounds in teff flour. The finding that the CFCS from B1.3 inhibited *S. cerevisiae* UCDFST 09-448 growth also suggests that B1.3 may be adapted to compete with yeast in teff injera. This result is consistent with the proximity of B1.3 to the olive-associated strains in the MLST phylogenetic tree. However, it is also noteworthy that B1.3 shares genetic similarity with the teff injera isolate (B1.1) and other grain-associated *L. plantarum* according to core genome comparisons.

Although disruptions in sucrose PTS systems may indicate why neither strain B1.3 nor 8.1 was able to grow in the presence of sucrose, the specific genes and pathways conferring the phenotypic variations observed in this study remain to be determined. To this regard, identification of the genome composition alone is insufficient to understand the full metabolic and functional potential of this species. For example, there still remains a lack of resolution in some PTS and other carbohydrate transport and metabolic pathways among lactobacilli (Gänzle and Follador, 2012; Zheng *et al*., 2015) and stress response mechanisms frequently involve numerous pathways with overlapping cell functions (e.g., membrane synthesis, protein turnover, and energy metabolism pathways) (Papadimitriou *et al*., 2016).

The genetic and phenotypic variation observed for the *L. plantarum* isolates indicate this species has evolved towards specialization in different plant-associated habitats (e.g., fruit vs cereal grains), but at the same time is under selective pressure for sustaining intraspecific diversity within those habitats, possibly as a mechanism promoting *L. plantarum* species stability through co-occurrence in those ecosystems (Maynard *et al*., 2019). Investigating this diversity and the importance of conserved and variable *L. plantarum* traits on plants and fermented plant foods is expected to be useful for understanding bacterial interactions and habitat partitioning in other complex host-associated (e.g., Lloyd-Price *et al*., 2017; Truong *et al*., 2017; Ma *et al*., 2020; Bongrand and Ruby, 2019) and environmental (e.g., Ellegaard *et al*., 2015; Props and Denef, 2020; Koch *et al*., 2020) sites wherein significant intraspecies diversity has been found but not yet understood. These findings may also be used to guide the selection of robust, multi-strain starter cultures that are suited to inter- and intra-species selection pressures in fruit and vegetable fermentations to result in optimal sensory and safety characteristics.

## Experimental Procedures

### Bacterial strains and growth conditions

*L. plantarum* strains used in this study are shown in **Table 1**. The isolates from olive fermentations and cactus fruit were described previously (Golomb *et al*., 2013; Tyler *et al*., 2016) and NCIMB8826R, a rifampicin-resistant variant (Yin *et al*., 2018) of strain NCIMB8826 (Hayward and Davis, 1956), was used as a reference. For *L. plantarum* isolation from injera batter, the batter was mixed with phosphate buffered saline (PBS, 137 mM NaCl, 2.7 mM KCl, 4.3 mM Na_2_HPO_4_-7H_2_O, 1.4 mM KH_2_PO_4_) (pH 7.2) at a ratio of 1:10. For isolation from boza and sourdough, the batter was mixed with physiological saline (145 mM NaCl) (pH 7.0) at a ratio of 1:10. For isolation from fermented tomatoes, three tomatoes were placed in sterile bags containing mesh filters (Nasco, Modesto, CA) with 1 ml of PBS (pH 7.2) and macerated by hand. Serial dilutions of the injera, boza, sourdough and tomato suspensions were then plated on de Man, Rogosa, and Sharpe (MRS) agar from a commercial source (BD, Franklin Lakes, NJ) (cMRS). Natamycin (25 μg/mL) (Dairy Connection, Wisconsin, WI) was included in the cMRS agar to inhibit fungal growth. The cMRS agar plates were incubated at 30 °C under aerobic or anaerobic conditions (BD BBL GasPak system (BD, Franklin Lakes, NJ) for 48 h. Single colony isolates were repeatedly streaked for isolation on cMRS prior to characterization. For phenotypic and genotypic analysis, the *L. plantarum* strains were routinely grown in cMRS without aeration at 30 °C.

### Strain identification and typing

*L. plantarum* 16S rRNA genes were amplified from individual colonies using the 27F and 1492R primers (Lane *et al*. 1991) (**Table S8**) with *ExTaq* DNA polymerase (TaKaRa, Shiga, Japan). Thermal cycling conditions were as follows: 95 °C for 3 min, 30 cycles of 94 °C for 30 sec, 50 °C for 30 sec, and 72 °C for 90 sec, and a final elongation step of 72 °C for 5 min. The PCR products were purified (Wizard SV Gel and PCR Clean-Up System (Promega, Madison, WI)) and sequenced at the UC Davis DNA Sequencing Facility http://dnaseq.ucdavis.edu/.The DNA sequences were compared against the National Center for Biotechnology Information (NCBI) database using the nucleotide Basic Local Alignment Search Tool (BLASTn) (https://blast.ncbi.nlm.nih.gov/Blast.cgi) and the Ribosomal Database Project (RDP) (http://rdp.cme.msu.edu/). Multiplex PCR targeting the *recA* gene was also used to confirm *L. plantarum* at the species level according to methods described by (Torriani *et al*., 2001) (**Table S8**). The 16S rRNA sequencing data for the strains in this study can be found National Center for Biotechnology Information (BankIt) under accession numbers MT937284-MT937296.

For multilocus sequence typing (MLST), genomic DNA was isolated with the Qiagen DNeasy Blood and Tissue Kit (Qiagen, Valencia, CA) according to the manufacturer’s instructions. PCR was then performed using primers targeting the variable regions of *L. plantarum pheS, pyrG, uvrC, recA, clpX, murC, groEL*, and *murE* (**Table S8**) (Xu *et al*., 2015). PCR amplification was preformed using *ExTaq* DNA polymerase (TaKaRa, Shiga, Japan) as previously described (Xu *et al*., 2015). The PCR products were sequenced in both directions using the forward and reverse primers at the UC Davis DNA Sequencing Facility (http://dnaseq.ucdavis.edu/) and Genewiz (South Plainfield, NJ). DNA sequences were aligned, trimmed, and analyzed using the MEGA 7.0 software package (Kumar *et al*., 2016). Based on the findings, unique nucleotide sequences for a gene were defined as an allele and unique allelic profiles were defined as a sequence type. The concatenate sequences in the order of *pheS, pyrG, uvrC, recA, clpX, murC, groEL*, and *murE* was used for phylogenetic tree analysis with maximum likelihood supported with a multilocus bootstrap approach using MEGA 7.0 (Kumar *et al*., 2016). For comparisons to other strains of *L. plantarum*, the sequences of 264 strains of *L. plantarum* were downloaded from the National Center for Biotechnology Information (NCBI) database (https://www.ncbi.nlm.nih.gov/), and a minimum spanning tree of the 278 strains was made using PHYLOVIZ Online (Ribeiro-Gonçalves *et al*., 2016). The MLST DNA sequences can be found in the National Center for Biotechnology Information (BankIt) under gene accession numbers MT864201-MT864291 and MT880889-MT880901,

### Genome sequencing, assembly, annotation, and analysis

Nine strains were selected for genome sequencing by either Illumina MiSeq (Illumina, San Diego, CA) (B1.1, WS1.1, 1B1, AJ11, BGM37, EL11) or Pacific Biosciences (PacBio, Menlo Park, CA) (B1.3, 8.1, K4) DNA sequencing methods. For the Illumina MiSeq, approximately 3×10^9^ cells were suspended in lysis buffer containing 200 mM NaCl, 20 mM EDTA, 500µl of 793 mM SDS and 300 mg of zirconium beads (0.1 mm, BioSpec Products, Bartlesville, OK). The cells were then mechanically lysed by bead-beating at 6.5m/s for 1 min with a FastPrep-24 (MP Biomedical, Santa Ana, CA). To obtain larger DNA fragments appropriate for PacBio DNA sequencing, total genomic DNA was extracted from each strain by incubating approximately 3×10^9^ cells in the presence of 20 mg/ml lysozyme (Sigma-Aldrich, St. Louis, MO) at 37 °C for 60 min. After extraction by either mechanical or enzymatic lysis, DNA was purified using phenol-chloroform and EtOH precipitation methods (Sambrook and Russell, 2006).

Illumina libraries were prepared for paired-end 250-bp sequencing (2 × 250 bp) using the Nextera DNA Flex Library kit (Illumina, San Diego, CA). The libraries were sequenced at the UC Davis Genome Center (Davis, CA) (https://genomecenter.ucdavis.edu/) on an Illumina MiSeq V2 according to the manufacturer’s protocol. Genomes were assembled with Spades (v3.12.0, using k-mers 31, 51, 71), and QUAST (v 4.6.3) was used to confirm assembly quality. The assembled genome sequences were then annotated with RASTtk and PATRIC (Wattam *et al*., 2017). PATRIC comprehensive genome analysis was run using default auto parameters. This program encompasses BayesHammer for read error correction, Velvet, IDBA, and Spades for assembly, and ARAST to verify assembly quality (Wattam *et al*., 2017). PacBio libraries were prepared and sequenced at the UC Davis Genome Center (Davis, CA) (https://genomecenter.ucdavis.edu/) on a Pacific Biosciences RSII instrument using P6-C4 sequencing chemistry. Sequence SMRTcell files were imported into the PacBio SMRT portal graphical interface unit (https://www.pacb.com/) for de novo assembly using the hierarchical genome-assembly process (HGAP) protocol (Chin *et al*., 2013) and RS HGAP Assembly 2 in Smart analysis version 2.3 software. The resulting assemblies were used for subsequent annotation with RASTtk (https://rast.nmpdr.org/) and PATRIC (Wattam *et al*., 2017). The whole genome sequencing data for this study can be found in the National Center for Biotechnology Information under the BioProject PRJNA598971.

EDGAR 2.0 was used the evaluate the size of the pangenome and identify the number of genes shared between all nine sequenced strains as well as to identify the phylogenetic relationships between the different strains (Blom *et al*., 2016). The pan and core genomes were identified, and the results were presented as ortholog sets. To evaluate phylogenetic relationships, concatenate core amino acid sequences were aligned using MUSCLE (Edgar, 2004). The resulting alignment was used to construct a phylogenetic tree using a maximum likelihood method with bootstrapping in MEGA 7.0 (Kumar *et al*., 2016). Anvi’o (v6.1) was used to group orthologous protein sequences into gene clusters for Cluster of Orthologues Group (COG) functional assignments using the program ‘anvi-pan-genome’ (Eren *et al*., 2015; Delmont and Eren, 2018) with the flags ‘-use-ncbi-blast’ (Altschul *et al*., 1990) and parameters ‘-minibit 0.5’ (Benedict *et al*., 2014) and ‘mcl-inflation 10’. COG frequency heat map with hierarchical clustering was generated using RStudio with the package ‘pheatmap’ (https://www.rstudio.com/). To confirm the truncation of *pts1BCA* in *L. plantarum* 8.1, the *pts1BCA* gene was amplified from genomic DNA from strains B1.3, K4, 8.1, and NCIMB8826R using the *pts1BCA_*trunF (5’-TCGTCACCGAGTGTTCGTTT) and *pts1BCA_*trunR (5’-AGTTGCTGGCCACTGTTCAT) primers (**Table S8**) and *ExTaq* DNA polymerase (TaKaRa, Shiga, Japan). Thermal cycling conditions were as follows: 95 °C for 3 min, 30 cycles of 94 °C for 30 sec, 50 °C for 30 sec, and 72 °C for 90 sec, and a final elongation step of 72 °C for 5 min. PCR products were visualized on a 1% agarose gel.

### Carbohydrate utilization

*L. plantarum* strains were first incubated in cMRS for 24 h at 30 °C. The cells were then collected by centrifugation at 5,000 x g for 5 min, washed twice in PBS to remove residual nutrients (pH 7.2), and then suspended in a modified MRS (mMRS) without beef extract or dextrose (pH 6.5) (De MAN *et al*., 1960). The cell suspensions were then distributed into 96-well microtiter plates (Thermo Fisher Scientific, Waltham, MA) at an optical density (OD) at 600 nm (OD_600_) of 0.2. To test the capacity to grow on different sugars, mMRS was amended to contain 2% (w/v) of D-glucose (111 mM) (Fisher Scientific, Fair Lawn, NJ), D-maltose monohydrate (55 mM) (Amresco, Solon, OH), sucrose (58 mM) (Sigma, St. Louis, MO), D-galactose (111 mM) (Fisher Scientific, Fair Lawn, NJ), D-raffinose pentahydrate (40 mM) (VWR International, Solon, OH), D-fructose (55 mM) (Fisher Scientific, Fair Lawn, NJ), D-xylose (133 mM) (Acros Organics, Morris Plains, NJ), D-ribose (133 mM) (Acros Organics, Morris Plains, NJ), or L-arabinose (133 mM) (Acros Organics, Morris Plains, NJ). The OD_600_ values were measured hourly for 48 h in a Synergy 2 microplate reader (Biotek, Winooski, VT) set at 30 °C without aeration.

### Growth during exposure to EtOH, SDS, NaCl, and pH 3.5

*L. plantarum* was incubated in cMRS for 24 h at 30 °C. The cells were then collected by centrifugation at 5,000 x g for 5 min, washed twice in PBS (pH 7.2), and then suspended in mMRS-glucose (2% (w/v) (111 mM) D-glucose) (pH 6.5). The cell suspensions were then distributed into 96-well microtiter plates containing mMRS-glucose amended to contain EtOH (8% (v/v) (174 mM) or 12% (v/v) (260 mM)), SDS (0.03% (w/v) (0.10 mM)), or NaCl (4% (w/v) (68 mM)). For measuring the effects of low pH, mMRS-glucose was adjusted to pH 3.5 with 1 M HCl. For measuring the effect of both low pH and high NaCl concentration, mMRS-glucose (pH 3.5) was supplemented with 4% (w/v) (68 mM) NaCl. Each strain was also incubated in mMRS diluted with water between (4 - 12% (v/v)) to control for dilution of mMRS due to amendment addition. The OD_600_ was used to monitor growth during incubation at 30 °C for 48 h without aeration using a Synergy 2 microplate reader (Biotek, Winooski, VT).

### Survival at pH 2 or 50 °C

For assessing acid tolerance, *L. plantarum* was incubated in cMRS for 24 h at 30 °C prior to collection by centrifugation at 5,000 x g for 5 min and washing twice in physiological saline (145 mM NaCl) (pH 7.0). *L. plantarum* was then inoculated at a concentration of 1 × 10^8^ cells/ml in physiological saline adjusted to pH 2 with 5 M HCl in 1.5mL tubes. Survival was measured after 0, 15, 30, and 60 min incubation at 30 °C. At each time point, three tubes were retrieved per stain for centrifugation at 10,000 x g for 1 min. The supernatant was discarded, and the resulting cell pellet was suspended in 1mL physiological saline (pH 7.0). Serial dilutions were then plated on cMRS agar and incubated at 30 °C for 48 h prior to colony enumeration.

### Survival at 50 °C

To measure thermal tolerance, *L. plantarum* was incubated in cMRS for 24 h at 30 °C prior to collection by centrifugation at 5,000 x g for 5 min and washing twice in PBS (pH 7.2). The suspensions were then distributed into 0.2 mL tubes at approximately 1 × 10^8^ CFU/ml and incubated in a C1000 Thermal Cycler (Bio-Rad Laboratories, Foster City, CA) at 50

°C for 0, 15, 30, and 60 min. At each time point, three tubes were retrieved per strain. Serial dilutions of the cell suspensions were plated onto cMRS agar and incubated at 30 °C for 48 h prior to colony enumeration.

### Biofilm formation assay

The potential for *L. plantarum* to form biofilms was assessed by measuring adherence to polystyrene according to previously described methods (Kopit *et al*., 2014) with several modifications. Briefly, 96-well polystyrene plates (Thermo Fisher Scientific, Waltham, MA) containing either mMRS-glucose, mMRS-fructose, or mMRS-sucrose were inoculated with *L. plantarum* to a starting OD_600_ of 0.2 and the plates were incubated at 30 °C for 48 h. The wells were then rinsed with PBS (pH 7.2), stained with 0.05% (w/v) crystal violet (CV), dried in an inverted position for 30 min, and then rinsed again three times with PBS (pH 7.2). Absorbance at OD_595_ was measured with a Synergy 2 microplate reader (Biotek, Winooski, VT) to determine adherence. Wells containing mMRS with the corresponding sugar without *L. plantarum* inoculum were included as controls.

### Yeast inhibition assay

*L. plantarum* cell-free culture supernatants (CFCS) were prepared from the spent media collected after *L. plantarum* incubation in cMRS for 24 h at 30 °C. CFCS was collected by centrifugation at 4,000 x g for 10 min at 4 °C followed by filtration of the supernatant through a 0.45 μm polyethersulfone (PES) filter (Genesee Scientific, San Diego, CA). To eliminate the effects of differences of pH on yeast inhibition, the CFCS was adjusted with lactic acid (1.3 M) to pH 3.8, the lowest pH reached by *L. plantarum* after incubation in cMRS (data not shown). *S. cerevisiae* UCDFST 09-448 (Golomb *et al*., 2013), a strain shown to cause olive tissue damage and spoilage during olive fermentations, was grown in Yeast Mold (YM) broth (BD, Franklin Lakes, NJ) for 24 h at 30 °C with aeration at 250 rpm. Cells were collected by centrifugation at 20,000 x g for 5 min at 4 °C and then washed twice with PBS. *S. cerevisiae* UCDFST 09-448 was then inoculated into 96-well microtiter plates containing 1:1 ratio of 2X YM and CFCS at a starting OD_600_ of 0.05. OD_600_ was measured in a Synergy 2 microplate reader (Biotek, Winooski, VT) set at 30 °C for 24 h aerated every hour by shaking for 10 sec before each read. Controls included *S. cerevisiae* UCDFST 09-448 incubated in YM and YM supplemented with cMRS (pH 3.8).

### Statistical analysis

Area under the curve (AUC) was used to examine the growth and survival of *L. plantarum* under different conditions (Sprouffske and Wagner, 2016). The AUC was calculated with GraphPad Prism 8 (Graph Pad Software, San Diego, CA). Hierarchical clustering was generated using RStudio with the package ‘pheatmap’ based on AUC values (https://www.rstudio.com/). Unpaired, two-tailed Student t-tests were used to compare between the different *L. plantarum* groups (e.g., brine- and grain-based fermentations). P values of <0.05 were considered significant.

## Acknowledgements

We would like to thank Nathan Lee and Brendan McCarthy-Sinclair for their assistance with conducting these assays. We would like to thank Menkir Tamrat for providing the injera from which we isolated *L. plantarum* B1.1 and B1.3. This work was supported by the USDA National Institute of Food and Agriculture (Grant No. CA-D-FST-2281-CG).

## References

Altschul, S.F., Gish, W., Miller, W., Myers, E.W., and Lipman, D.J. (1990) Basic local alignment search tool. J Mol Biol 215: 403–410.

Aquilanti, L., Santarelli, S., Silvestri, G., Osimani, A., Petruzzelli, A., and Clementi, F. (2007) The microbial ecology of a typical Italian salami during its natural fermentation. Int J of Food Microbiol 120: 136–145.

Barache, N., Ladjouzi, R., Belguesmia, Y., Bendali, F., and Drider, D. (2020) Abundance of Lactobacillus plantarum strains with beneficial attributes in blackberries (Rubus sp.), fresh figs (Ficus carica), and prickly pears (Opuntia ficus-indica) grown and harvested in Algeria. Probiotics & Antimicro Prot.

Benedict, M.N., Henriksen, J.R., Metcalf, W.W., Whitaker, R.J., and Price, N.D. (2014) ITEP: An integrated toolkit for exploration of microbial pan-genomes. BMC Genomics 15: 8.

Blom, J., Kreis, J., Spänig, S., Juhre, T., Bertelli, C., Ernst, C., and Goesmann, A. (2016) EDGAR 2.0: an enhanced software platform for comparative gene content analyses. Nucleic Acids Res 44: W22–W28.

Bolnick, D.I., Amarasekare, P., Araújo, M.S., Bürger, R., Levine, J.M., Novak, M., et al. (2011) Why intraspecific trait variation matters in community ecology. Trends Eco Evol 26: 183–192.

Bongrand, C. and Ruby, E.G. (2019) Achieving a multi-strain symbiosis: strain behavior and infection dynamics. ISME J 13: 698–706.

Bravo-Ferrada, B.M., Gonçalves, S., Semorile, L., Santos, N.C., Tymczyszyn, E.E., and Hollmann, A. (2015) Study of surface damage on cell envelope assessed by AFM and flow cytometry of Lactobacillus plantarum exposed to ethanol and dehydration. J Appl Microbio 118: 1409–1417.

Brizuela, N., Tymczyszyn, E.E., Semorile, L.C., Valdes La Hens, D., Delfederico, L., Hollmann, A., and Bravo-Ferrada, B. (2019) Lactobacillus plantarum as a malolactic starter culture in winemaking: A new (old) player? Electron J Biotech 38: 10–18.

Cataldi, T.R.I., Margiotta, G., Iasi, L., Di Chio, B., Xiloyannis, C., and Bufo, S.A. (2000) Determination of sugar compounds in olive plant extracts by anion-exchange chromatography with pulsed amperometric detection. Anal Chem 72: 3902–3907.

Chin, C.-S., Alexander, D.H., Marks, P., Klammer, A.A., Drake, J., Heiner, C., et al. (2013) Nonhybrid, finished microbial genome assemblies from long-read SMRT sequencing data. Nat Methods 10: 563–569.

Choi, S., Jin, G.-D., Park, J., You, I., and Kim, E.B. (2018) Pan-genomics of Lactobacillus plantarum revealed group-specific genomic profiles without habitat association. J Microbiol Biotechnol 28: 1352–1359.

Ciocia, F., McSweeney, P.L.H., Piraino, P., and Parente, E. (2013) Use of dairy and non-dairy Lactobacillus plantarum, Lactobacillus paraplantarum and Lactobacillus pentosus strains as adjuncts in cheddar cheese. Dairy Sci & Technol 93: 623–640.

Corcoran, B.M., Stanton, C., Fitzgerald, G., and Ross, R.P. (2008) Life under stress: the probiotic stress response and how it may be manipulated. Curr Pharm Des 14: 1382–1399.

Crakes, K.R., Rocha, C.S., Grishina, I., Hirao, L.A., Napoli, E., Gaulke, C.A., et al. (2019) PPARα-targeted mitochondrial bioenergetics mediate repair of intestinal barriers at the host–microbe intersection during SIV infection. PNAS 116: 24819–24829.

De MAN, J.C., Rogosa, M., and Sharpe, M.E. (1960) A medium for the cultivation of Lactobacilli. Journal of Applied Bacteriology 23: 130–135.

Delgado, S., Flórez, A.B., and Mayo, B. (2005) Antibiotic Susceptibility of Lactobacillus and Bifidobacterium species from the human gastrointestinal tract. Curr Microbiol 50: 202–207.

Delmont, T.O. and Eren, A.M. (2018) Linking pangenomes and metagenomes: the Prochlorococcus metapangenome. PeerJ 6: e4320.

Di Cagno, R., Surico, R.F., Siragusa, S., De Angelis, M., Paradiso, A., Minervini, F., et al. (2008) Selection and use of autochthonous mixed starter for lactic acid fermentation of carrots, French beans or marrows. Int J Food Microbiol 127: 220–228.

Duar, R.M., Lin, X.B., Zheng, J., Martino, M.E., Grenier, T., Pérez-Muñoz, M.E., et al. (2017) Lifestyles in transition: evolution and natural history of the genus Lactobacillus. FEMS Microbiol Rev 41: S27–S48.

Edgar, R.C. (2004) MUSCLE: multiple sequence alignment with high accuracy and high throughput. Nucleic Acids Res 32: 1792–1797.

Ehlers, B.K., Damgaard, C.F., and Laroche, F. (2016) Intraspecific genetic variation and species coexistence in plant communities. Biol Lett 12: 20150853.

Ellegaard, K.M., Tamarit, D., Javelind, E., Olofsson, T.C., Andersson, S.G., and Vásquez, A. (2015) Extensive intra-phylotype diversity in lactobacilli and bifidobacteria from the honeybee gut. BMC Genomics 16: 284.

Endo, A., Maeno, S., Tanizawa, Y., Kneifel, W., Arita, M., Dicks, L., and Salminen, S. (2018) Fructophilic lactic acid bacteria, a unique group of fructose-fermenting microbes. Appl Environ Microbiol 84: e01290–18.

Ercolini, D. (2017) Exciting strain-level resolution studies of the food microbiome. Microb Biotechnol 10: 54–56.

Eren, A.M., Esen, Ö.C., Quince, C., Vineis, J.H., Morrison, H.G., Sogin, M.L., and Delmont, T.O. (2015) Anvi’o: an advanced analysis and visualization platform for ‘omics data. PeerJ 3: e1319.

Fernández Ramírez, M.D., Smid, E.J., Abee, T., and Nierop Groot, M.N. (2015) Characterization of biofilms formed by Lactobacillus plantarum WCFS1 and food spoilage isolates. Int J Food Microbio 207: 23–29.

Ferrando, V., Quiberoni, A., Reinheimer, J., and Suárez, V. (2016) Functional properties of Lactobacillus plantarum strains: A study in vitro of heat stress influence. Food Microbiol 54: 154–161.

Ferrando, V., Quiberoni, A., Reinhemer, J., and Suárez, V. (2015) Resistance of functional Lactobacillus plantarum strains against food stress conditions. Food Microbiol 48: 63–71.

Filannino, P., Cardinali, G., Rizzello, C.G., Buchin, S., Angelis, M.D., Gobbetti, M., and Cagno, R.D. (2014) Metabolic responses of Lactobacillus plantarum strains during fermentation and storage of vegetable and fruit juices. Appl Environ Microbiol 80: 2206–2215.

France, M.T., Mendes-Soares, H., and Forney, L.J. (2016) Genomic comparisons of Lactobacillus crispatus and Lactobacillus iners reveal potential ecological drivers of community composition in the vagina. Appl Environ Microbiol 82: 7063–7073.

Gänzle, M.G. and Follador, R. (2012) Metabolism of oligosaccharides and starch in Lactobacilli: A Review. Front Microbiol 3: 340.

Gheziel, C., Russo, P., Arena, M.P., Spano, G., Ouzari, H.-I., Kheroua, O., et al. (2019) Evaluating the probiotic potential of Lactobacillus plantarum strains from Algerian infant feces: towards the design of probiotic starter cultures tailored for developing countries. Probiotics & Antimicro Prot 11: 113–123.

Golomb, B.L., Morales, V., Jung, A., Yau, B., Boundy-Mills, K.L., and Marco, M.L. (2013) Effects of pectinolytic yeast on the microbial composition and spoilage of olive fermentations. Food Microbiol 33: 97–106.

Gómez-González, S., Ruiz-Jiménez, J., Priego-Capote, F., and Luque de Castro, M.D. (2010) Qualitative and quantitative sugar orofiling in olive fruits, leaves, and stems by gas chromatography-tandem mass spectrometry (GC-MS/MS) after ultrasound-assisted leaching. J Agric Food Chem 58: 12292–12299.

Görke, B. and Stülke, J. (2008) Carbon catabolite repression in bacteria: many ways to make the most out of nutrients. Nat Rev Microbiol 6: 613–624.

van de Guchte, M., Penaud, S., Grimaldi, C., Barbe, V., Bryson, K., Nicolas, P., et al. (2006) The complete genome sequence of Lactobacillus bulgaricus reveals extensive and ongoing reductive evolution. Proc Natl Acad Sci U S A 103: 9274–9279.

Guidone, A., Zotta, T., Ross, R.P., Stanton, C., Rea, M.C., Parente, E., and Ricciardi, A. (2014) Functional properties of Lactobacillus plantarum strains: A multivariate screening study. LWT - Food Sci and Technol 56: 69–76.

Hayward, A.C. and Davis, G.H.G. (1956) The isolation and classification of Lactobacillus strains from Italian saliva samples. Br Dent J 101: 2733–2741.

Hurtado, A., Reguant, C., Bordons, A., and Rozès, N. (2012) Lactic acid bacteria from fermented table olives. Food Microbiol 31: 1–8.

Jose, N.M., Bunt, C.R., and Hussain, M.A. (2015) Comparison of microbiological and probiotic characteristics of Lactobacilli isolates from dairy food products and animal rumen contents. Microorganisms 3: 198–212.

Koch, H., Germscheid, N., Freese, H.M., Noriega-Ortega, B., Lücking, D., Berger, M., et al. (2020) Genomic, metabolic and phenotypic variability shapes ecological differentiation and intraspecies interactions of Alteromonas macleodii. Sci Rep 10: 809.

Kopit, L.M., Kim, E.B., Siezen, R.J., Harris, L.J., and Marco, M.L. (2014) Safety of the surrogate microorganism Enterococcus faecium NRRL B-2354 for use in thermal process validation. Appl Environ Microbiol 80: 1899–1909.

Kremling, A., Geiselmann, J., Ropers, D., and de Jong, H. (2015) Understanding carbon catabolite repression in Escherichia coli using quantitative models. Trends in Microbiol 23: 99–109.

Kumar, S., Stecher, G., and Tamura, K. (2016) MEGA7: Molecular Evolutionary Genetics Analysis Version 7.0 for bigger datasets. Mol Biol Evol 33: 1870–1874.

Lloyd-Price, J., Mahurkar, A., Rahnavard, G., Crabtree, J., Orvis, J., Hall, A.B., et al. (2017) Strains, functions and dynamics in the expanded Human Microbiome Project. Nature 550: 61–66.

Ma, B., France, M.T., Crabtree, J., Holm, J.B., Humphrys, M.S., Brotman, R.M., and Ravel, J. (2020) A comprehensive non-redundant gene catalog reveals extensive within-community intraspecies diversity in the human vagina. Nature Commun 11: 1–13.

Marco, M. (2010) Lactobacillus plantarum in foods. In Encyclopedia of Biotechnology in Agriculture and Food. CRC Press, pp. 360–362.

Martino, M.E., Bayjanov, J.R., Caffrey, B.E., Wels, M., Joncour, P., Hughes, S., et al. (2016) Nomadic lifestyle of Lactobacillus plantarum revealed by comparative genomics of 54 strains isolated from different habitats. Environ Microbiol 18: 4974–4989.

Maynard, D.S., Serván, C.A., Capitán, J.A., and Allesina, S. (2019) Phenotypic variability promotes diversity and stability in competitive communities. Ecology Letters 22: 1776–1786.

Miller, E.R., Kearns, P.J., Niccum, B.A., O’Mara Schwartz, J., Ornstein, A., and Wolfe, B.E. (2019) Establishment Limitation constrains the abundance of lactic acid bacteria in the Napa cabbage phyllosphere. Appl Environ Microbiol 85: e00269–19.

Mills, S., Stanton, C., Fitzgerald, G.F., and Ross, R.P. (2011) Enhancing the stress responses of probiotics for a lifestyle from gut to product and back again. Microb Cell Fact 10: S19.

Molenaar, D., Bringel, F., Schuren, F.H., Vos, W.M. de, Siezen, R.J., and Kleerebezem, M. (2005) Exploring Lactobacillus plantarum genome diversity by using microarrays. J Bacteriol 187: 6119–6127.

Mukhopadhyay, A. (2015) Tolerance engineering in bacteria for the production of advanced biofuels and chemicals. Trends in Microbiol 23: 498–508.

Papadimitriou, K., Alegría, Á., Bron, P.A., Angelis, M. de, Gobbetti, M., Kleerebezem, M., et al. (2016) Stress physiology of lactic acid bacteria. Microbiol Mol Biol Rev 80: 837–890.

Parente, E., Ciocia, F., Ricciardi, A., Zotta, T., Felis, G.E., and Torriani, S. (2010) Diversity of stress tolerance in Lactobacillus plantarum, Lactobacillus pentosus and Lactobacillus paraplantarum: A multivariate screening study. Int J Food Microbiol 144: 270–279.

Parichehreh, S., Tahmasbi, G., Sarafrazi, A., Imani, S., and Tajabadi, N. (2018) Isolation and identification of Lactobacillus bacteria found in the gastrointestinal tract of the dwarf honeybee, Apis florea Fabricius, 1973 (Hymenoptera: Apidae). Apidologie 49: 430–438.

Prete, R., Long, S.L., Joyce, S.A., and Corsetti, A. (2020) Genotypic and phenotypic characterization of food-associated Lactobacillus plantarum isolates for potential probiotic activities. FEMS Microbiol Lett.

Props, R. and Denef, V.J. (2020) Temperature and nutrient levels correspond with lineage-specific microdiversification in the ubiquitous and abundant freshwater genus Limnohabitans. Appl Environ Microbiol 86: e00140–20.

Ribeiro-Gonçalves, B., Francisco, A.P., Vaz, C., Ramirez, M., and Carriço, J.A. (2016) PHYLOViZ Online: web-based tool for visualization, phylogenetic inference, analysis and sharing of minimum spanning trees. Nucleic Acids Res 44: W246–W251.

Salvetti, E., Harris, H.M.B., Felis, G.E., and O’Toole, P.W. (2018) Comparative genomics of the genus Lactobacillus reveals robust phylogroups that provide the basis for reclassification. Appl Environ Microbiol 84: e00993–18.

Sambrook, J. and Russell, D.W. (2006) Purification of Nucleic Acids by Extraction with Phenol:Chloroform. Cold Spring Harb Protoc 2006: pdb.prot4455.

Saulnier, D.M.A., Molenaar, D., Vos, W.M. de, Gibson, G.R., and Kolida, S. (2007) Identification of prebiotic fructooligosaccharide metabolism in Lactobacillus plantarum WCFS1 through microarrays. Appl Environ Microbiol 73: 1753–1765.

Seddik, H.A., Bendali, F., Gancel, F., Fliss, I., Spano, G., and Drider, D. (2017) Lactobacillus plantarum and its probiotic and food potentialities. Probiotics & Antimicro Prot 9: 111–122.

Seddon, A.M., Curnow, P., and Booth, P.J. (2004) Membrane proteins, lipids and detergents: not just a soap opera. Biochimica et Biophysica Acta (BBA) - Biomembranes 1666: 105–117.

Siezen, R.J. and van Hylckama Vlieg, J.E. (2011) Genomic diversity and versatility of Lactobacillus plantarum, a natural metabolic engineer. Microb Cell Fact 10: S3.

Siezen, R.J., Tzeneva, V.A., Castioni, A., Wels, M., Phan, H.T.K., Rademaker, J.L.W., et al. (2010) Phenotypic and genomic diversity of Lactobacillus plantarum strains isolated from various environmental niches. Environ Microbiol 12: 758–773.

Siragusa, S., De Angelis, M., Calasso, M., Campanella, D., Minervini, F., Di Cagno, R., and Gobbetti, M. (2014) Fermentation and proteome profiles of Lactobacillus plantarum strains during growth under food-like conditions. J Proteom 96: 366–380.

Sprouffske, K. and Wagner, A. (2016) Growthcurver: an R package for obtaining interpretable metrics from microbial growth curves. BMC Bioinformatics 17: 172.

Succi, M., Pannella, G., Tremonte, P., Tipaldi, L., Coppola, R., Iorizzo, M., et al. (2017) Sub-optimal pH preadaptation improves the survival of Lactobacillus plantarum strains and the malic acid consumption in wine-like medium. Front Microbiol 8: 470.

Sun, Z., Harris, H.M.B., McCann, A., Guo, C., Argimón, S., Zhang, W., et al. (2015) Expanding the biotechnology potential of lactobacilli through comparative genomics of 213 strains and associated genera. Nat Commun 6: 8322.

Torriani, S., Felis, G.E., and Dellaglio, F. (2001) Differentiation of Lactobacillus plantarum, L. pentosus, and L. paraplantarum by recA gene sequence analysis and multiplex PCR assay with recA gene-derived primers. Appl Environ Microbiol 67: 3450–3454.

Truong, D.T., Tett, A., Pasolli, E., Huttenhower, C., and Segata, N. (2017) Microbial strain-level population structure and genetic diversity from metagenomes. Genome Res 27: 626–638.

Tyler, C. a., Kopit, L., Doyle, C., Yu, A. o., Hugenholtz, J., and Marco, M. l. (2016) Polyol production during heterofermentative growth of the plant isolate Lactobacillus florum 2F. J Appl Microbiol 120: 1336–1345.

Veen, H. van B. de, Abee, T., Tempelaars, M., Bron, P.A., Kleerebezem, M., and Marco, M.L. (2011) Short-and long-term adaptation to ethanol stress and its cross-protective consequences in Lactobacillus plantarum. Appl Environ Microbiol 77: 5247–5256.

Wattam, A.R., Davis, J.J., Assaf, R., Boisvert, S., Brettin, T., Bun, C., et al. (2017) Improvements to PATRIC, the all-bacterial bioinformatics database and analysis Resource Center. Nucleic Acids Res 45: D535–D542.

Westby, A., Nuraida, L., Owens, J.D., and Gibbs, P.A. (1993) Inability of Lactobacillus plantarum and other lactic acid bacteria to grow on D-ribose as sole source of fermentable carbohydrate. J Appl Bacteriol 75: 168–175.

Xu, H., Sun, Z., Liu, W., Yu, J., Song, Y., Lv, Q., et al. (2014) Multilocus sequence typing of Lactococcus lactis from naturally fermented milk foods in ethnic minority areas of China. J Dairy Sci 97: 2633–2645.

Yang, J., Cao, Y., Cai, Y., and Terada, F. (2010) Natural populations of lactic acid bacteria isolated from vegetable residues and silage fermentation. J Dairy Sci 93: 3136–3145.

Yin, W., Wang, Y., Liu, L., and He, J. (2019) Biofilms: The microbial “protective clothing” in extreme environments. Int J Mol Sci 20: 3423.

Yin, X., Heeney, D.D., Srisengfa, Y.T., Chen, S.-Y., Slupsky, C.M., and Marco, M.L. (2018) Sucrose metabolism alters Lactobacillus plantarum survival and interactions with the microbiota in the digestive tract. FEMS Microbiol Ecol 94: fiy084.

Yu, A.O., Leveau, J.H.J., and Marco, M.L. (2020) Abundance, diversity and plant-specific adaptations of plant-associated lactic acid bacteria. Environ Microbiol Rep 12: 16–29.

Zago, M., Scaltriti, E., Bonvini, B., Fornasari, M.E., Penna, G., Massimiliano, L., et al. (2017) Genomic diversity and immunomodulatory activity of Lactobacillus plantarum isolated from dairy products. Benef Microbes 8: 597–604.

Zaragoza, J., Bendiks, Z., Tyler, C., Kable, M.E., Williams, T.R., Luchkovska, Y., et al. (2017) Effects of exogenous yeast and bacteria on the microbial population dynamics and outcomes of olive fermentations. mSphere 2: e00315–16.

Zheng, J., Ruan, L., Sun, M., and Gänzle, M. (2015) A genomic view of Lactobacilli and Pediococci demonstrates that phylogeny matches ecology and physiology. Appl Environ Microbiol 81: 7233–7243.

Zheng, J., Wittouck, S., Salvetti, E., Franz, C.M.A.P., Harris, H.M.B., Mattarelli, P., et al. (2020) A taxonomic note on the genus Lactobacillus: Description of 23 novel genera, emended description of the genus Lactobacillus Beijerinck 1901, and union of Lactobacillaceae and Leuconostocaceae. Int J Syst Evol Microbiol 70: 2782–2858.

