## Supplementary Materials for "Strain diversity of plant-associated *Lactiplantibacillus plantarum*"

**Table S1. Allelic profiles of *L. plantarum* strains according to MLST[a].**

| Strain | *clpX* | *groEL* | *murC* | *murE* | *pheS* | *pyrG* | *recA* | *uvrC* | ST[b] |
|---|---|---|---|---|---|---|---|---|---|
| **AJ11** | 2 | 2 | 2 | 2 | 2 | 2 | 2 | 2 | **1** |
| **BGM55** | 1 | 1 | 1 | 1 | 1 | 1 | 1 | 1 | **2** |
| **BGM37** | 5 | 5 | 6 | 1 | 5 | 7 | 6 | 1 | **3** |
| **BGM40** | 7 | 6 | 7 | 2 | 7 | 10 | 8 | 1 | **4** |
| **EL11** | 3 | 4 | 4 | 4 | 3 | 4 | 4 | 1 | **5** |
| **K4** | 2 | 2 | 3 | 1 | 6 | 9 | 3 | 4 | **6** |
| **8.1** | 2 | 3 | 3 | 3 | 2 | 3 | 3 | 1 | **7** |
| **W1.1** | 2 | 2 | 5 | 5 | 4 | 5 | 2 | 3 | **8** |
| **B1.1** | 2 | 2 | 3 | 2 | 4 | 5 | 7 | 3 | **9** |
| **B1.3** | 2 | 7 | 3 | 7 | 9 | 12 | 3 | 6 | **10** |
| **T2.5** | 9 | 2 | 3 | 3 | 2 | 8 | 3 | 5 | **11** |
| **WS1.1** | 6 | 2 | 3 | 3 | 2 | 8 | 3 | 3 | **12** |
| **1B1** | 8 | 2 | 3 | 1 | 8 | 11 | 3 | 3 | **13** |
| **NCIMB8826R** | 4 | 2 | 5 | 6 | 2 | 6 | 5 | 3 | **14** |

[a] MLST based on (Xu *et al.*, 2015)
[b] Sequence Type (ST).

**Table S2. Average AUC values of *L. plantarum* under different growth conditions.**

| | **Carbon Source [a]** | | | | | | | | | **Environmental Stress [b]** | | | | | |
|---|---|---|---|---|---|---|---|---|---|---|---|---|---|---|---|
| **Strain** | **Glucose[b]** | **Maltose[b]** | **Sucrose[b]** | **Raffinose[b]** | **Galactose[b]** | **Fructose[b]** | **Ribose[b]** | **Arabinose[b]** | **Xylose[b]** | **8% EtOH[c]** | **12% EtOH[c]** | **0.03% SDS[c]** | **pH 3.5[c]** | **4% NaCl[c]** | **pH 3.5 + 4% NaCl[c]** |
| **AJ11** | 138 | 143 | 137 | 114 | 103 | 117 | 36 | 25 | 24 | 115 | 113 | 106 | 58 | 114 | 15 |
| **BGM55** | 137 | 139 | 133 | 116 | 138 | 116 | 22 | 13 | 16 | 108 | 100 | 100 | 60 | 122 | 13 |
| **BGM37** | 145 | 148 | 143 | 121 | 148 | 137 | 40 | 41 | 15 | 140 | 83 | 120 | 67 | 119 | 19 |
| **BGM40** | 140 | 142 | 140 | 115 | 101 | 120 | 33 | 19 | 16 | 105 | 101 | 110 | 56 | 116 | 18 |
| **EL11** | 136 | 139 | 134 | 105 | 106 | 100 | 17 | 15 | 16 | 118 | 108 | 103 | 42 | 108 | 10 |
| **K4** | 134 | 136 | 138 | 125 | 123 | 114 | 16 | 14 | 18 | 102 | 98 | 112 | 46 | 101 | 14 |
| **8.1** | 135 | 140 | 46 | 38 | 91 | 76 | 22 | 13 | 18 | 122 | 60 | 110 | 30 | 117 | 12 |
| **W1.1** | 140 | 134 | 134 | 118 | 117 | 98 | 17 | 12 | 16 | 106 | 55 | 92 | 29 | 103 | 11 |
| **B1.1** | 140 | 144 | 138 | 56 | 125 | 121 | 25 | 14 | 19 | 122 | 82 | 104 | 37 | 93 | 11 |
| **B1.3** | 91 | 54 | 34 | 92 | 98 | 113 | 31 | 12 | 18 | 40 | 11 | 107 | 32 | 37 | 12 |
| **T2.5** | 135 | 140 | 130 | 60 | 114 | 102 | 21 | 16 | 17 | 102 | 56 | 113 | 48 | 110 | 13 |
| **WS1.1** | 137 | 144 | 129 | 68 | 117 | 108 | 23 | 13 | 23 | 104 | 102 | 114 | 49 | 116 | 14 |
| **1B1** | 140 | 143 | 138 | 118 | 119 | 123 | 17 | 15 | 18 | 130 | 121 | 120 | 48 | 107 | 16 |
| **NCIMB8826R** | 132 | 137 | 129 | 113 | 120 | 112 | 33 | 28 | 21 | 102 | 73 | 106 | 54 | 106 | 13 |

[a] Average AUC of three independent replicates after incubation for 48 h at 30 °C in MRS (pH 6.5) without beef extract or dextrose was supplemented with 2% (w/v) of each of the following carbohydrate sources: glucose, maltose, sucrose, raffinose, galactose, fructose, ribose, arabinose, and xylose

[b] Average AUC of three independent replicates after incubation in mMRS-glucose supplemented with 8% (v/v) EtOH, 8% (v/v) EtOH and then 12% (v/v) EtOH (12% EtOH), 0.03% (w/v) SDS, 4% (w/v) NaCl or set at pH 3.5 without or with 4% (w/v) NaCl.

**Table S3. Final $OD_{600}$ of *L. plantarum* under different growth conditions.**

| | Carbon Source [a] | | | | | | | | | Environmental Stress [b] | | | | | |
|---|---|---|---|---|---|---|---|---|---|---|---|---|---|---|---|
| **Strain** | **Glucose** | **Maltose** | **Sucrose** | **Raffinose** | **Galactose** | **Fructose** | **Ribose** | **Arabinose** | **Xylose** | **8% EtOH** | **12% EtOH** | **0.03% SDS** | **pH 3.5** | **4% NaCl** | **pH 3.5 + 4% NaCl** |
| **AJ11** | 3.33[a] | 3.39 | 3.35 | 3.08 | 2.63 | 2.98 | 1.36 | 0.97 | 0.53 | 3.11 | 3.26 | 2.79 | 2.28 | 3.00 | 0.40 |
| **BGM55** | 3.31 | 3.41 | 3.27 | 3.13 | 3.41 | 3.05 | 1.46 | 0.30 | 0.31 | 2.94 | 2.96 | 2.52 | 2.37 | 3.20 | 0.26 |
| **BGM37** | 3.53 | 3.62 | 3.52 | 3.38 | 3.62 | 3.44 | 2.93 | 2.99 | 0.26 | 4.02 | 2.55 | 3.05 | 2.61 | 3.16 | 0.52 |
| **BGM40** | 3.37 | 3.46 | 3.44 | 3.08 | 2.58 | 3.06 | 2.03 | 0.48 | 0.29 | 2.83 | 3.05 | 2.85 | 2.27 | 3.07 | 0.50 |
| **EL11** | 3.30 | 3.45 | 3.33 | 3.09 | 2.93 | 2.64 | 0.40 | 0.43 | 0.32 | 3.55 | 3.24 | 2.62 | 1.73 | 3.20 | 0.22 |
| **K4** | 3.36 | 3.35 | 3.50 | 3.26 | 3.13 | 2.99 | 0.33 | 0.3 | 0.40 | 2.86 | 2.93 | 2.91 | 1.98 | 2.96 | 0.39 |
| **8.1** | 3.29 | 3.34 | 1.04 | 2.54 | 2.67 | 1.83 | 0.58 | 0.25 | 0.39 | 3.39 | 2.46 | 2.80 | 1.49 | 3.22 | 0.31 |
| **W1.1** | 3.53 | 3.52 | 3.51 | 3.44 | 3.54 | 2.69 | 0.38 | 0.25 | 0.32 | 3.49 | 2.87 | 2.60 | 1.26 | 3.21 | 0.23 |
| **B1.1** | 3.58 | 3.60 | 3.56 | 3.33 | 3.52 | 3.31 | 0.70 | 0.31 | 0.45 | 3.59 | 3.56 | 3.06 | 1.61 | 3.23 | 0.26 |
| **B1.3** | 2.93 | 1.95 | 0.95 | 2.65 | 3.08 | 3.13 | 0.90 | 0.24 | 0.37 | 1.91 | 0.23 | 2.94 | 1.15 | 1.96 | 0.27 |
| **T2.5** | 3.33 | 3.47 | 2.98 | 3.07 | 3.15 | 2.63 | 0.50 | 0.35 | 0.36 | 2.94 | 3.51 | 2.93 | 2.06 | 2.98 | 0.38 |
| **WS1.1** | 3.37 | 3.49 | 2.99 | 3.14 | 3.17 | 2.77 | 0.58 | 0.25 | 0.58 | 2.99 | 3.37 | 2.98 | 2.07 | 3.14 | 0.45 |
| **1B1** | 3.44 | 3.51 | 3.41 | 3.38 | 3.25 | 3.19 | 0.39 | 0.29 | 0.36 | 3.69 | 3.77 | 3.07 | 1.97 | 2.94 | 0.48 |
| **NCIMB8826R** | 3.25 | 3.39 | 3.21 | 3.06 | 3.08 | 2.91 | 1.51 | 1.94 | 0.45 | 2.81 | 2.38 | 2.71 | 2.20 | 2.98 | 0.32 |

[a] Average final $OD_{600}$ of three independent replicates after incubation for 48 h at 30 °C in MRS (pH 6.5) without beef extract or dextrose was supplemented with 2% (w/v) of each of the following carbohydrate sources: glucose, maltose, sucrose, raffinose, galactose, fructose, ribose, arabinose, and xylose

[b] Average final $OD_{600}$ of three independent replicates after incubation in mMRS-glucose supplemented with 8% (v/v) EtOH, 8% (v/v) EtOH and then 12% (v/v) EtOH (12% EtOH), 0.03% (w/v) SDS, 4% (w/v) NaCl or set at pH 3.5 without or with 4% (w/v) NaCl.

**Table S4. Growth rates of *L. plantarum* in mMRS containing different sugars.**

| Strain | Glucose | Maltose | Sucrose | Raffinose | Galactose | Fructose | Ribose | Arabinose | Xylose |
|---|---|---|---|---|---|---|---|---|---|
| **AJ11** | 0.44[a] | 0.41 | 0.42 | 0.28 | 0.24 | 0.39 | 0.14 | 0.12 | 0.14 |
| **BGM55** | 0.43 | 0.39 | 0.40 | 0.24 | 0.38 | 0.39 | 0.12 | 0.12 | 0.10 |
| **BGM37** | 0.45 | 0.42 | 0.42 | 0.20 | 0.42 | 0.36 | 0.10 | 0.07 | 0.10 |
| **BGM40** | 0.45 | 0.41 | 0.43 | 0.29 | 0.25 | 0.39 | 0.12 | 0.12 | 0.10 |
| **EL11** | 0.42 | 0.40 | 0.40 | 0.16 | 0.22 | 0.34 | 0.10 | 0.10 | 0.11 |
| **K4** | 0.40 | 0.38 | 0.36 | 0.41 | 0.28 | 0.35 | 0.09 | 0.03 | 0.09 |
| **8.1** | 0.42 | 0.40 | 0.32 | 0.28 | 0.21 | 0.38 | 0.10 | 0.06 | 0.11 |
| **W1.1** | 0.39 | 0.31 | 0.31 | 0.29 | 0.18 | 0.32 | 0.11 | 0.05 | 0.10 |
| **B1.1** | 0.38 | 0.36 | 0.36 | 0.15 | 0.23 | 0.29 | 0.10 | 0.04 | 0.10 |
| **B1.3** | 0.20 | 0.15 | 0.20 | 0.12 | 0.16 | 0.27 | 0.12 | 0.02 | 0.08 |
| **T2.5** | 0.43 | 0.39 | 0.39 | 0.16 | 0.21 | 0.38 | 0.11 | 0.09 | 0.10 |
| **WS1.1** | 0.44 | 0.39 | 0.40 | 0.19 | 0.21 | 0.36 | 0.12 | 0.06 | 0.12 |
| **1B1** | 0.45 | 0.42 | 0.43 | 0.17 | 0.27 | 0.41 | 0.08 | 0.07 | 0.11 |
| **NCIMB8826R** | 0.43 | 0.39 | 0.40 | 0.27 | 0.31 | 0.38 | 0.15 | 0.12 | 0.11 |

Average growth rates of three independent replicates are provided ($h^{-1}$) after incubation in mMRS supplemented with 2% (w/v) of each of the following carbohydrate sources: glucose, maltose, sucrose, raffinose, galactose, fructose, ribose, arabinose, or xylose.

**Table S5. Growth rates of *L. plantarum* exposed to different environmental stresses.**

| **Strain** | **8% EtOH** | **12% EtOH** | **0.03% SDS** | **pH 3.5** | **4% NaCl** | **pH 3.5 + 4% NaCl** |
|---|---|---|---|---|---|---|
| **AJ11** | 0.28[a] | 0.14 | 0.31 | 0.09 | 0.36 | 0.02 |
| **BGM55** | 0.30 | 0.10 | 0.30 | 0.10 | 0.36 | 0.02 |
| **BGM37** | 0.32 | 0.10 | 0.35 | 0.12 | 0.38 | 0.04 |
| **BGM40** | 0.34 | 0.10 | 0.31 | 0.08 | 0.34 | 0.02 |
| **EL11** | 0.33 | 0.11 | 0.30 | 0.07 | 0.27 | 0.01 |
| **K4** | 0.25 | 0.09 | 0.34 | 0.09 | 0.31 | 0.01 |
| **8.1** | 0.30 | 0.11 | 0.38 | 0.07 | 0.34 | 0.01 |
| **W1.1** | 0.22 | 0.10 | 0.22 | 0.06 | 0.25 | 0.01 |
| **B1.1** | 0.25 | 0.10 | 0.25 | 0.07 | 0.28 | 0.01 |
| **B1.3** | 0.09 | NA | 0.32 | 0.05 | 0.06 | NA |
| **T2.5** | 0.25 | 0.10 | 0.34 | 0.09 | 0.35 | 0.01 |
| **WS1.1** | 0.24 | 0.12 | 0.35 | 0.07 | 0.34 | 0.02 |
| **1B1** | 0.31 | 0.11 | 0.36 | 0.10 | 0.38 | 0.03 |
| **NCIMB8826R** | 0.27 | 0.11 | 0.31 | 0.11 | 0.32 | 0.02 |

Average growth rates of three independent replicates after incubation in mMRS-glucose supplemented with 8% (v/v) EtOH, 8% (v/v) EtOH and then 12% (v/v) EtOH (12% EtOH), 0.03% (w/v) SDS, 4% (w/v) NaCl or set at pH 3.5 without or with 4% (w/v) NaCl.
An "NA" indicates that the strain did not grow under this condition

**Table S3. Growth characteristics of *S. cerevisiae* UCDFST 09-448 incubated in *L. plantarum* cell free supernatant (CFCS).**

| CFCS | Growth rate ($h^{-1}$) | Final $OD_{600}$ | AUC |
|---|---|---|---|
| **AJ11** | **0.30 ± 0.007** | **1.64 ± 0.05** | **31.64 ± 0.29** |
| **BGM55** | **0.29 ± 0.03[b]** | **1.57 ± 0.16** | **31.66 ± 1.31** |
| **BGM37** | **0.30 ± 0.006** | **1.60 ± 0.11** | **32.33 ± 0.61** |
| **BGM40** | **0.30 ± 0.008** | **1.56 ± 0.11** | **31.22 ± 0.61** |
| **EL11** | **0.31 ± 0.003** | **1.65 ± 0.08** | **32.58 ± 1.08** |
| **K4** | **0.31 ± 0.007** | **1.79 ± 0.04** | **33.94 ± 0.23** |
| **8.1** | **0.30 ± 0.007** | **1.64 ± 0.05** | **32.05 ± 0.27** |
| **W1.1** | **0.32 ± 0.005** | **1.72 ± 0.02** | **33.21 ± 0.12** |
| **B1.1** | **0.31 ± 0.007** | **1.74 ± 0.03** | **33.69 ± 0.13** |
| **B1.3** | **0.30 ± 0.007** | **1.77 ± 0.07** | **33.76 ± 1.06** |
| **T2.5** | **0.29 ± 0.003** | **1.51 ± 0.03** | **29.71 ± 0.25** |
| **WS1.1** | **0.29 ± 0.002** | **1.51 ± 0.04** | **29.54 ± 0.19** |
| **1B1** | **0.31 ± 0.003** | **1.64 ± 0.005** | **32.02 ± 0.03** |
| **NCIMB8826R** | **0.29 ± 0.01** | **1.57 ± 0.08** | **31.20 ± 0.46** |
| cMRS (pH 3.8) | 0.35 ± 0.004 | 2.20 ± 0.05 | 40.59 ± 0.26 |

Avg ± stdev of three independent replicates was determined after 24 h incubation at 30 °C. Bold type indicates significantly different ($p < 0.05$) growth characteristics compared to growth in cMRS (pH 3.8) controls using unpaired, two-tailed Student T-test ($p < 0.05$).

**Table S7. Distribution of gene clusters across *L. plantarum* genomes based on COG categories.**

| | AJ11 | BGM37 | EL11 | K4 | 8.1 | B1.1 | B1.3 | WS1.1 | 1B1 |
|---|---|---|---|---|---|---|---|---|---|
| **C: Energy production and conversion** | 124 | 118 | 119 | 115 | 117 | 118 | 111 | 121 | 118 |
| **D: Cell Division and Chromosome Partitioning** | 40 | 45 | 37 | 37 | 36 | 40 | 37 | 41 | 41 |
| **E: Amino acid metabolism and transport** | 198 | 203 | 192 | 191 | 198 | 192 | 181 | 209 | 193 |
| **F: Nucleotide metabolism and transport** | 89 | 93 | 92 | 89 | 98 | 89 | 91 | 98 | 88 |
| **G: Carbohydrate metabolism and transport** | 238 | 256 | 239 | 216 | 228 | 211 | 206 | 222 | 230 |
| **H: Coenzyme metabolism** | 111 | 104 | 101 | 99 | 104 | 114 | 108 | 108 | 102 |
| **I: Lipid metabolism** | 73 | 76 | 73 | 72 | 74 | 75 | 71 | 76 | 69 |
| **J: Translation** | 198 | 203 | 198 | 199 | 202 | 198 | 201 | 202 | 201 |
| **K: Transcription** | 276 | 291 | 274 | 258 | 267 | 244 | 215 | 279 | 268 |
| **L: Replication, recombination and repair** | 106 | 125 | 111 | 118 | 142 | 116 | 117 | 139 | 126 |
| **M: Cell wall structure and biogenesis and outer membrane** | 167 | 152 | 153 | 135 | 148 | 162 | 148 | 160 | 159 |
| **N: Secretion, motility and chemotaxis** | 17 | 22 | 22 | 24 | 20 | 20 | 22 | 22 | 18 |
| **O: Molecular chaperones and related functions** | 87 | 89 | 88 | 86 | 96 | 84 | 83 | 91 | 89 |
| **P: Inorganic ion transport and metabolism** | 114 | 118 | 120 | 116 | 124 | 109 | 104 | 139 | 117 |
| **Q: Secondary metabolites biosynthesis, transport, and catabolism** | 33 | 37 | 32 | 28 | 35 | 31 | 32 | 33 | 32 |
| **R: General functional prediction** | 205 | 206 | 206 | 194 | 199 | 192 | 176 | 203 | 199 |
| **S: No functional prediction** | 193 | 190 | 190 | 182 | 192 | 183 | 171 | 200 | 193 |
| **T: Signal Transduction** | 104 | 110 | 109 | 107 | 105 | 101 | 102 | 110 | 108 |
| **U: Intracellular trafficking, secretion, and vesicular transport** | 16 | 14 | 14 | 16 | 18 | 17 | 14 | 18 | 18 |
| **V: Defense mechanisms** | 76 | 85 | 85 | 81 | 87 | 78 | 79 | 95 | 80 |
| **X: Mobilome: prophages, transposons** | 49 | 86 | 63 | 88 | 179 | 117 | 352 | 132 | 104 |

Functional COG categories were assigned to gene clusters using Anvi'o (v6.1) (Eren et al., 2015).

**Table S8. PCR primers used in this study.**

| Primer Name | Primer Sequence (5' → 3') | References |
|---|---|---|
| **27F** | AGAGTTTGATCMTGGCTCAG | Lane *et al.* 1991 |
| **1492R** | GGTTACCTTGTTACGACTT | Lane *et al.* 1991 |
| **paraF** | GTCACAGGCATTACGAAAAC | Torriani *et al.* 2001 |
| **pentF** | CAGTGGCGCGGTTGATATC | Torriani *et al.* 2001 |
| **planF** | CCGTTTATGCGGAACACCTA | Torriani *et al.* 2001 |
| **pREV** | TCGGGATTACCAAACATCAC | Torriani *et al.* 2001 |
| ***pheS*_primerF** | CCGTGAAGAACTGGAACA | Xu *et al.* 2015 |
| ***pheS*_primerR** | CCTAACCCAAAGGCAAAA | Xu et al. 2015 |
| ***pyrG*_primerF** | AGTGATTTAGGTTCCGACAA | Xu *et al.* 2015 |
| ***pyrG*_primerR** | TGCATTCCCAAGCAGATA | Xu *et al.* 2015 |
| ***uvrC*_primerF** | GATCATTTATGTGGGTAAGGC | Xu *et al.* 2015 |
| ***uvrC*_primerR** | TGACACTACTGGGAACAAGC | Xu *et al.* 2015 |
| ***recA*_primerF** | TTTTAGTTGTTGACTCGGTGGC | Xu *et al.* 2015 |
| ***recA*_primerR** | TTCCGCTGGTGTCGCTTT | Xu *et al.* 2015 |
| ***clpX*_primerF** | ATCGCCAAGAAGAGTGAA | Xu *et al.* 2015 |
| ***clpX*_primerR** | ATAATCGAGCGTAGACCC | Xu *et al.* 2015 |
| ***murC*_primerF** | TATCGCTCCCACCAGTTA | Xu *et al.* 2015 |
| ***murC*_primerR** | CGGCCAAGATTTCCTTAT | Xu *et al.* 2015 |
| ***groEL*_primerF** | CGGCTACTTATCACAATACA | Xu *et al.* 2015 |
| ***groEL*_primerR** | GCCTTCTAAACCAGCATT | Xu *et al.* 2015 |
| ***murE*_primerF** | ACTAATAAGGTCGCTGTTCTG | Xu *et al.* 2015 |
| ***murE*_primerR** | TTTAGCGGCTTCTTCACT | Xu *et al.* 2015 |
| ***pts1BCA*_trunF** | TCGTCACCGAGTGTTCGTTT | This study |
| ***pts1BCA*_trunR** | AGTTGCTGGCCACTGTTCAT | This study |

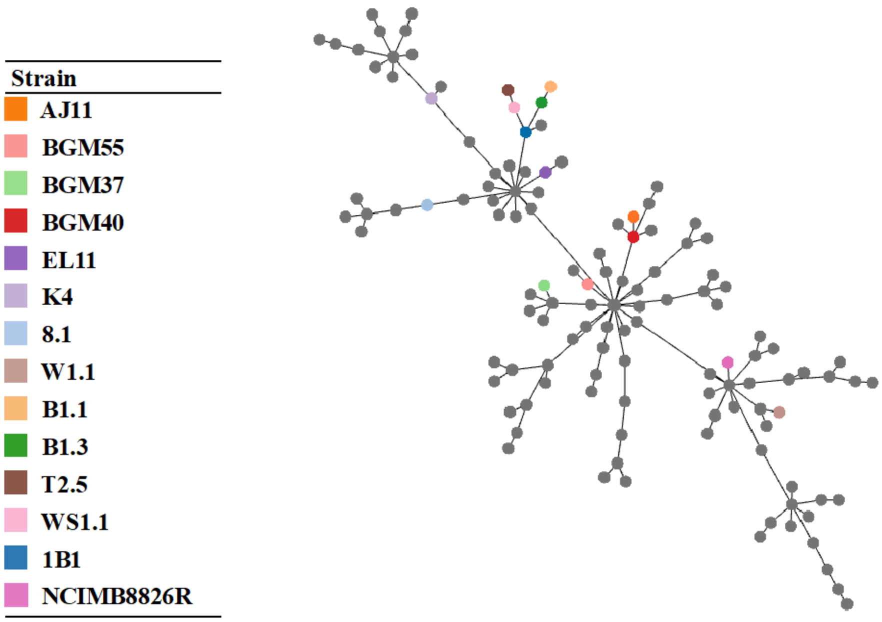


**Fig. S1. Minimum spanning tree of *L. plantarum* using MLST.** The MLST profile was determined from 264 strains using a multi-locus typing scheme developed previously (Xu et al. 2015). A minimum spanning tree was made using PHYLOVIZ Online (https://online.phyloviz.net/index). Strains with a common ST were grouped into a node. Nodes were colored based on the presence of the referenced strain within that node.

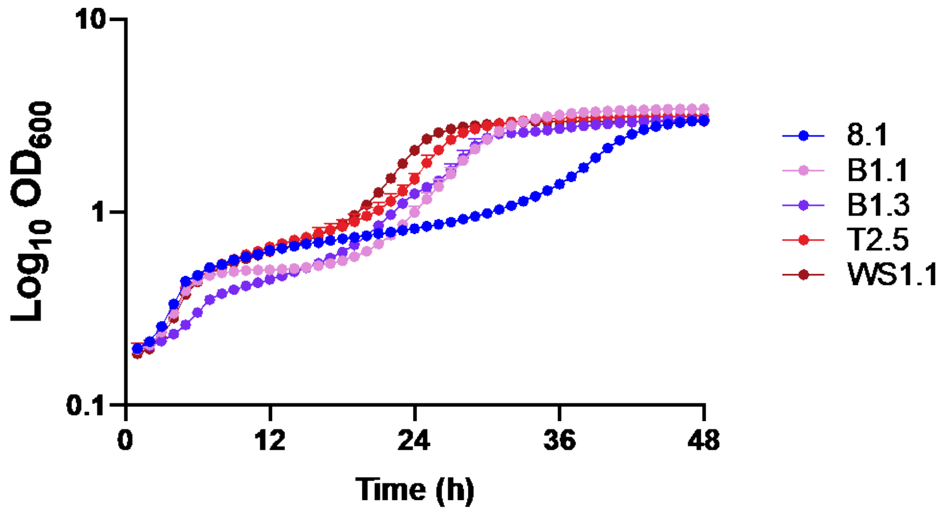


**Fig S2. Growth of *L. plantarum* in mMRS containing 2% (w/v) raffinose after repeated incubation in that culture medium.** *L. plantarum* grown overnight in mMRS-raffinose for 24 h was inoculated into mMRS-raffinose and incubated at 30 °C for 48 h. The average AUC values improved for all strains except for B1.3. Compared to AUC values measured after direct inoculation in cMRS for 24 h, the average AUC increased with successive passage for strain 8.1 from 38 to 58, strain B1.1 from 56 to 83, strain T2.5 from 60 to 88 and strain WS1.1 from 68 to 89 or decreased for strain B1.3 from 92 to 76. The avg ± stdev $OD_{600}$ of three replicates wells are shown.

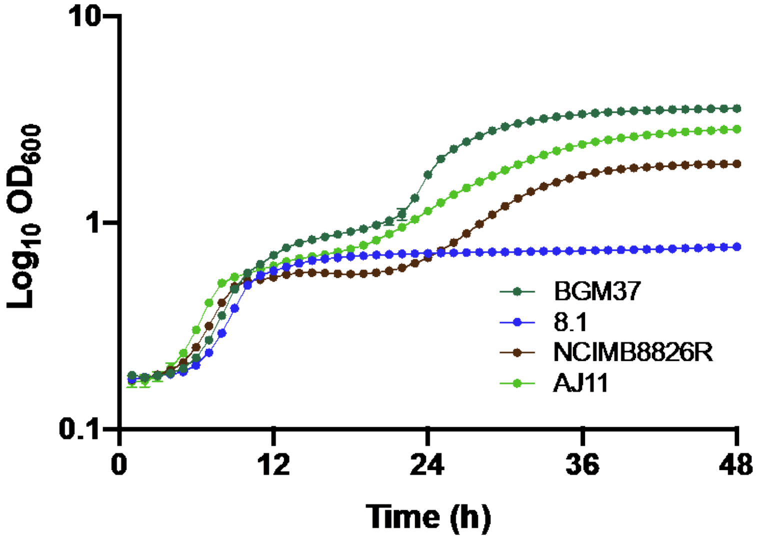


**Fig S3. Growth of *L. plantarum* in mMRS containing 2% (w/v) ribose after repeated-incubation in that culture medium.** *L. plantarum* grown overnight in mMRS containing 2% (w/v) ribose was subsequently inoculated again into mMRS containing 2% (w/v) ribose incubated at 30 °C for 48 h. Compared to AUC values measured after direct inoculation in cMRS for 24 h, the average AUC (n = 3) increased with successive passage for strain BGM37 from 40 to 94, strain NCIMB8826R from 33 to 49, strain AJ11 from 36 to 70, however, the AUC did not change for strain 8.1 (22). The avg ± stdev $OD_{600}$ of three replicates wells are shown.

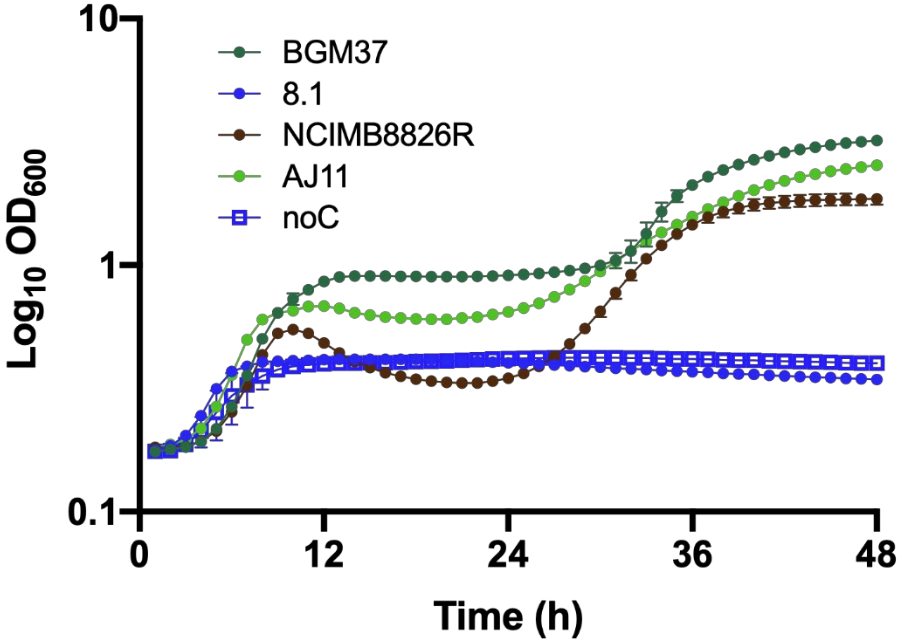


**Fig S4. Growth of *L. plantarum* in mMRS containing 2% (w/v) arabinose after repeated-incubation in that culture medium.** *L. plantarum* grown overnight in mMRS containing 2% (w/v) arabinose was subsequently inoculated again into mMRS containing 2% (w/v) arabinose and incubated at 30˚C °C for 48 h. The avg ± se $OD_{600}$ of three replicates wells are shown. The average (n = 3) AUC improved for all strains, except strain 8.1. Compared to AUC values measured after direct inoculation in cMRS for 24 h, the average AUC (n = 3) increased with successive passage for strain BGM37 from 41 to 66, strain NCIMB8826R from 28 to 40, strain AJ11 from 25 to 53, however, the AUC did not change for strain 8.1 (13). A representative strain (8.1) grown in mMRS without added 2% (w/v) raffinose was included as a reference.

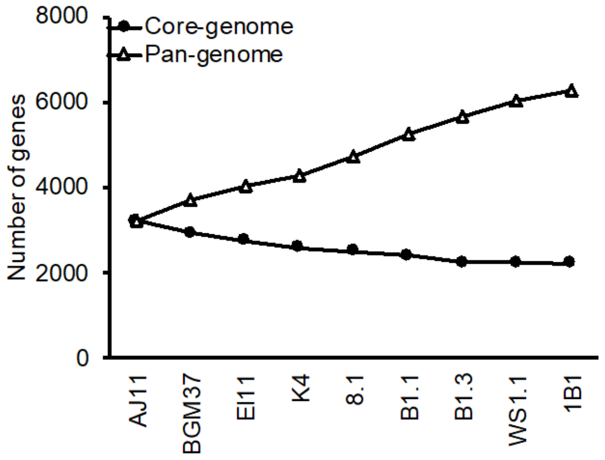


**Fig S5. The core-genome and pan-genome of nine *L. plantarum* strains examined in this study.** The estimated sizes of the *L. plantarum* core genome (●) pangenome (Δ) are shown as a cumulative function of the strains compared.
